# Insights into structure of *Penicillium funiculosum* LPMO and its synergistic saccharification performance with CBH1 on high substrate loading upon simultaneous overexpression

**DOI:** 10.1101/2020.04.16.045914

**Authors:** Olusola A. Ogunyewo, Anmoldeep Randhawa, Mayank Gupta, Vemula Chandra Kaladhar, Praveen Kumar Verma, Syed Shams Yazdani

## Abstract

Lytic polysaccharide monooxygenases (LPMOs) are crucial industrial enzymes required in the biorefinery industry as well as in natural carbon cycle. These enzymes known to possess auxiliary activity are produced by numerous bacterial and fungal species to assist in the degradation of cellulosic biomass. In this study, we annotated and performed structural analysis of an uncharacterized thermostable LPMO from *Penicillium funiculosum* (PfLPMO9) in an attempt to understand nature of this enzyme in biomass degradation. PfLPMO9 exhibited 75% and 36% structural identity to *Thermoascus aurantiacus* (TaLPMO9A) and *Lentinus similis* (LsLPMO9A), respectively. Analysis of the molecular interactions during substrate binding revealed that PfLPMO9 demonstrated a higher binding affinity with a ΔG free energy of -46 k kcal/mol when compared with that of TaLPMO9A (−31 kcal/mol). The enzyme was further found to be highly thermostable at elevated temperature with a half-life of ∼88 h at 50 °C. Furthermore, multiple fungal genetic manipulation tools were employed to simultaneously overexpress this LPMO and Cellobiohydrolase I (CBH1) in catabolite derepressed strain of *Penicillium funiculosum, Pf*Mig1^88^, in order to improve its saccharification performance towards acid pretreated wheat straw (PWS) at 20% substrate loading. The resulting transformants showed ∼200% and ∼66% increase in LPMO and Avicelase activities, respectively. While the secretomes of individually overexpressed LPMO and CBH1-strains increased saccharification of PWS by 6% and 13%, respectively, over *Pf*Mig1^88^ at same enzyme concentration, the simultaneous overexpression of these two genes led to 20% increase in saccharification efficiency over *Pf*Mig1^88^, which accounted for 82% saccharification of PWS at 20% substrate loading.

**Importance:** Enzymatic hydrolysis of cellulosic biomass by cellulases continues to be a significant bottleneck in the development of second-generation bio-based industries. While efforts are being intensified at how best to obtain indigenous cellulase for biomass hydrolysis, the high production cost of this enzyme remains a crucial challenge confronting its wide availability for efficient utilization of cellulosic materials. This is because it is challenging to get an enzymatic cocktail with balanced activity from a single host. This report provides for the first time the annotation and structural analysis of an uncharacterized thermostable lytic polysaccharide monooxygenase (LPMO) gene in *Penicillium funiculosum* and its impact in biomass deconstruction upon overexpression in catabolite derepressed strain of *P. funiculosum*. Cellobiohydrolase I (CBH1) which is the most important enzyme produced by many cellulolytic fungi for saccharification of crystalline cellulose was further overexpressed simultaneously with the LPMO. The resulting secretome was analyzed for enhanced LPMO and exocellulase activities with the corresponding improvement in its saccharification performance at high substrate loading by ∼20% using a minimal amount of protein.

## Introduction

Lignocellulosic biomass continues to attract worldwide attention as a cheaper source of energy generation owing to its relative abundance, renewable nature, and its wide availability in the environment as a low-cost substrate. This lignocellulosic biomass, which consists of cellulose, hemicellulose and lignin, is a potential renewable resource that can be hydrolyzed and converted to fermentable sugars for the production of bioethanol for renewable energy generation (1–3). Cellulases, which are crucial industrial enzymes required in biofuel generation, are mainly produced in abundance by filamentous fungi. They are vital for cellulose degradation by catalyzing the hydrolysis of β-1,4 glycosidic bonds in cellulosic materials, to produce short cellulo-oligosaccharides and glucose (4). These enzymes from filamentous fungi perform a central role in the global carbon cycle by degrading insoluble cellulose to soluble sugars (5–7). Over time, the secretomes of many fungal genera such as *Trichoderma, Aspergillus, Neurospora* and *Penicillium* have been studied for their ability to produce cellulolytic enzymes. Although, the secretome from *T. reesei* which serves as a model fungus for cellulase production is widely used in commercial cocktails; complete saccharification of cellulosic biomass by its secretome is usually not achievable. This is because the secretome of *T. reesei* under cellulase-inducing conditions secretes mainly cellobiohydrolases (about 85%) while other cellulase components such as β-glucosidase and endoglucanase which participate in biomass hydrolysis are secreted in a lower amount and have to be produced in other fungal sources to complement the cellobiohydrolase in commercial cocktails (8–10).

Furthermore, while there is increasing attention on how to fortify the secretome of many cellulase producing strains for better biomass hydrolysis, recent efforts have identified the significant roles of some synergistic or auxiliary proteins in promoting cellulase activity. Effective utilization of these synergistic proteins such as expansins, swollenin, hydrophobins, lytic polysaccharide monooxygenase (LPMO) and so many more have been reported to significantly reduce the total cellulase loading needed to achieve about 80 – 90 % cellulose conversion required by industries for bioethanol production (11–14). LPMOs, a family of recently discovered auxiliary proteins are unique in terms of their capability of catalyzing the cleavage of glycosidic bonds of polysaccharides via an oxidative mechanism rather than hydrolytic means. The mechanism of cellulose cleavage by LPMOs involve the reduction of Cu^2+^ at the active site by some redox partners. This consequently brings about an attack on the pyranose ring of the glucose moieties at the C-1 or C-4 position, thereby destabilizing the adjacent glycosidic bond and breaking it by an elimination reaction (15, 16). These LPMOs targets the crystalline regions of the cellulose surface which are typically more recalcitrant to cellulase action and introduce nicks on the substrate surface thereby providing extra chain ends for glycoside hydrolases (GHs) to act (17). This action subsequently improves the reaction kinetics as well as enhance enzymatic synergy during saccharification (17, 18). However, the critical drawback of not having a single strain which could secrete a balanced ratio of all the cellulase component (both hydrolytic and accessory) subsequently affects the overall enzyme cost required for biomass saccharification especially at high substrate concentration. This challenge has, therefore, prompted many kinds of research into looking for better strains which could secrete to a large extent, all the key enzyme components required for effective biomass hydrolysis.

In our search for an indigenous producer of cellulolytic enzymes, we identified *Penicillium funiculosum* (NCIM1228), a filamentous fungus whose secretome exhibited an outstanding potential for hydrolyzing cellulosic biomass (19). The proteomic analysis of its secretome revealed that about 58% of the total proteins secreted under cellulase inducing conditions belongs to the Carbohydrate-Active Enzymes (CAZymes) family. When the ratio of the cellobiohydrolases (CBH I and II) that are the most important and abundant protein present in the secretome of any cellulase producing strain was assessed, it was found that the proportion of CBHs in the secretome of *P. funiculosum* was only 15% of the total proteins which is contrary to the secretome of *T. reesei* where the CBHs represented 95% of the total proteins (19, 20). Furthermore, during the structure-function characterization of CBH1 enzyme of *P. funiculosum*, it was found that CBH1 of *P. funiculosum* can hydrolyse crystalline biomass with about five-fold higher efficiency than its counterpart from well-known industrial host *Trichoderma reesei* (21). The enhancement in enzymatic activity was found to be mainly caused by the catalytic domain of PfCBH1 when the subdomains of PfCBH1 and TrCBH1 were swapped (22). Interestingly, further proteomic analysis of the secretome under cellulase inducing condition also revealed the detection of the unique oxidative enzyme; LPMO (formerly GH61) belonging to the AA9 family, which was also confirmed by enzyme assay (19). It was hypothesized that the presence of this LPMO in the secretome could have contributed to the higher saccharification property of the secretome of this fungus when compared with cellulase from other fungal hosts. (15, 23, 24).

Therefore, to further improve this strain as an industrial workhorse for production of lignocellulolytic enzymes, the regulatory pathway was first manipulated genetically for the deregulation of cellulolytic genes in its genome (25). This deregulation was done by deleting a homolog of a carbon catabolite repressor (CCR) Mig1 in *Penicillium funiculosum* NCIM1228 thereby yielding a catabolite derepressed strain, *Penicillium funiculosum* Mig1^88^ (*Pf*Mig1^88^), which showed more than two-fold improvement in enzyme activity and protein titer in the secretome (25). When the saccharification potential of *Pf*Mig1^88^ secretome was evaluated towards different chemically pretreated cellulosic biomass, it was found that *Pf*Mig1^88^ secretome could hydrolyze ∼75% of total cellulose into glucose from nitric acid + ammonium hydroxide pretreated biomass in 72 h at 30 mg/g DBW (dry biomass weight) and 20% biomass loading (26). Under this pretreatment condition, even without recombinant expression or supplementation, *Pf*Mig1^88^ secretome could also achieve the benchmark of ∼70% hydrolysis of sugars using 20 mg/g DBW at 20% solid loading recommended for cellulolytic enzymes to be utilized for second-generation biofuels (27). However, pretreatment with either nitric acid or ammonium hydroxide alone resulted in the relatively lower release of 60% and 63% sugars, respectively, which suggested that accessibility to the cellulosic fibres was hindered by lignin and hemicellulose (26). This drawback thus necessitated the need to prospect into better ways of increasing the cellulase content of the secretome to increase the probability of substrate-enzyme proximity even in the presence of lignin and hemicellulose.

Based on the above, this study was designed to further improve the efficiency of the secretome of *Pf*Mig1^88^ strain so as to enhance its saccharification performance towards highly recalcitrant biomass at higher substrate loading with a minimal amount of protein. This was done by fortifying the secretome of *Pf*Mig1^88^ through the engineering of the key hydrolytic and oxidative enzymes which participates in saccharification of crystalline cellulosic biomass. Analysis of the genome sequence of *P. funiculosum* NCIM1228 has established that the fungal strain has only a single copy of gene coding for LPMO, and only a single copy of gene coding for CBH1, while the same fungal strain has more than one copy of genes coding for other cellulase components such as β-glucosidase, endoglucanase and xylanase (19, 28). This, therefore, motivated the idea to see if the overexpression of any of these two genes (LPMO and CBH1) could positively improve the efficiency of this secretome towards highly recalcitrant biomass. Considering that the structures of lytic polysaccharide monooxygenase (LPMO) from *P. funiculosum* have never been reported, we first performed *in-silico* structural modelling and thermostability study to understand novel features of this enzyme. We then overexpressed the two selected genes in the catabolite derepressed strain of *P. funiculosum* independently and simultaneously. Secretome from each of the resulting transformants was then assessed for improved production of cellulolytic enzymes as well as their saccharification efficiency towards acid pretreated wheat straw at 20% substrate loading.

## Results and Discussion

### Phylogenetic distribution and computational analysis of LPMO gene in *Penicillium funiculosum*

From different studies and reports, it is known that the number of LPMO genes in fungi varies and can range from one single gene to as many as twelve depending on the organism (29–33). Therefore, for us to ascertain the number of LPMO genes present in *P. funiculosum*, we first performed whole-genome annotation for this organism. The annotation indicated the presence of a single gene for LPMO (Fig. 1A, Supplementary file S1), which belongs to the AA9 family in the CAZy database and thus designated as PfLPMO9. The preliminary analysis of the PfLPMO9 nucleotide and the encoded protein sequence showed that it consists of 1020 bp, including two introns, which encodes 310 amino acids with a theoretical molecular weight of 32 kDa (Fig. 1A). Since there was limited report on LPMO from *P. funiculosum*, we proceeded further to perform the phylogenetic analysis with the LPMO sequences available at NCBI database to assess its neighbours. We found LPMO ortholog from *Talaromyces pinophilus and Penicillium occitanis* in the tree being closest sharing the same branch, while the ones from well-studied fungi, such as *Trichoderma ressei*, and *Neurospora crassa* were at different branches (Fig. 1B). The phylogenetic analysis places PfLPMO9 in a cluster together with C1/C4-oxidizing LPMOs which are active on cello-oligosaccharides suggesting that this LPMO could mediate oxidative cleavage of cellulosic biomass at either the reducing (C1) or non-reducing (C4) positions (15). This was in agreement with a recent report on LPMO from a mutated strain of *P. funiculosum* which was able to mediate cellulose oxidation at the C1 end while there was no available information about its C4 oxidation (34). To assess putative structural differences (if any) between the C1/C4 – oxidizing LPMOs, a structural model of PfLPMO9 was built and used for comparison with already characterized LPMOs. Since there are no 3-D structures available for LPMO from *P. funiculosum* in public database, we first retrieved from PDB the full-length amino acid sequences corresponding to the catalytic domains of the C1/C4 oxidizing LPMOs whose 3-D structures were available and subjected to multiple sequence alignment (MSA) alongside with the catalytic domain of PfLPMO9. Keeping in mind that proteins with same evolutionary relationships do have a certain percentage of their amino acid residues conserved, we found a reasonable degree of amino acid conservation between the various LPMO catalytic domains retrieved and analyzed (Fig. 2). The three residues His22, His107 and Tyr196, which are involved in copper coordination and binding of cello-oligosaccharides chains were seen to be conserved among all the LPMOs analyzed. The alignment showed that PfLPMO9 exhibited 75% identity to that of *Thermoascus aurantiacus* (TaLPMO9A) while 53%, 50% and 36% identity was seen with LPMOs from *Neurospora crassa* (NCU07760), *Trichoderma ressei* (HjLPMO9B) and *Lentinus similis* (LsAA9A) respectively. A homology model of PfLPMO9 was further constructed using protein references mentioned in the Materials and Methods section (Fig. S1A). The modelled structure was evaluated for accuracy by Ramachandran plot validation using PROCHECK module of the PDBSum server. An array of the Φ and Ψ distributions of the non-glycine, non-proline residues on the Ramachandran plot are summarized in Fig. S1B. The Ramachandran statistics revealed that 90.5% of amino acid residues from the modelled structure were incorporated in the favoured regions (A, B, and L) of the plot while 8.9% of the residues were in the allowed regions (a, b, l, and p) of the plot, which suggests that the model is reasonable and reliable for homologous structural analysis.

**Fig. 1:**
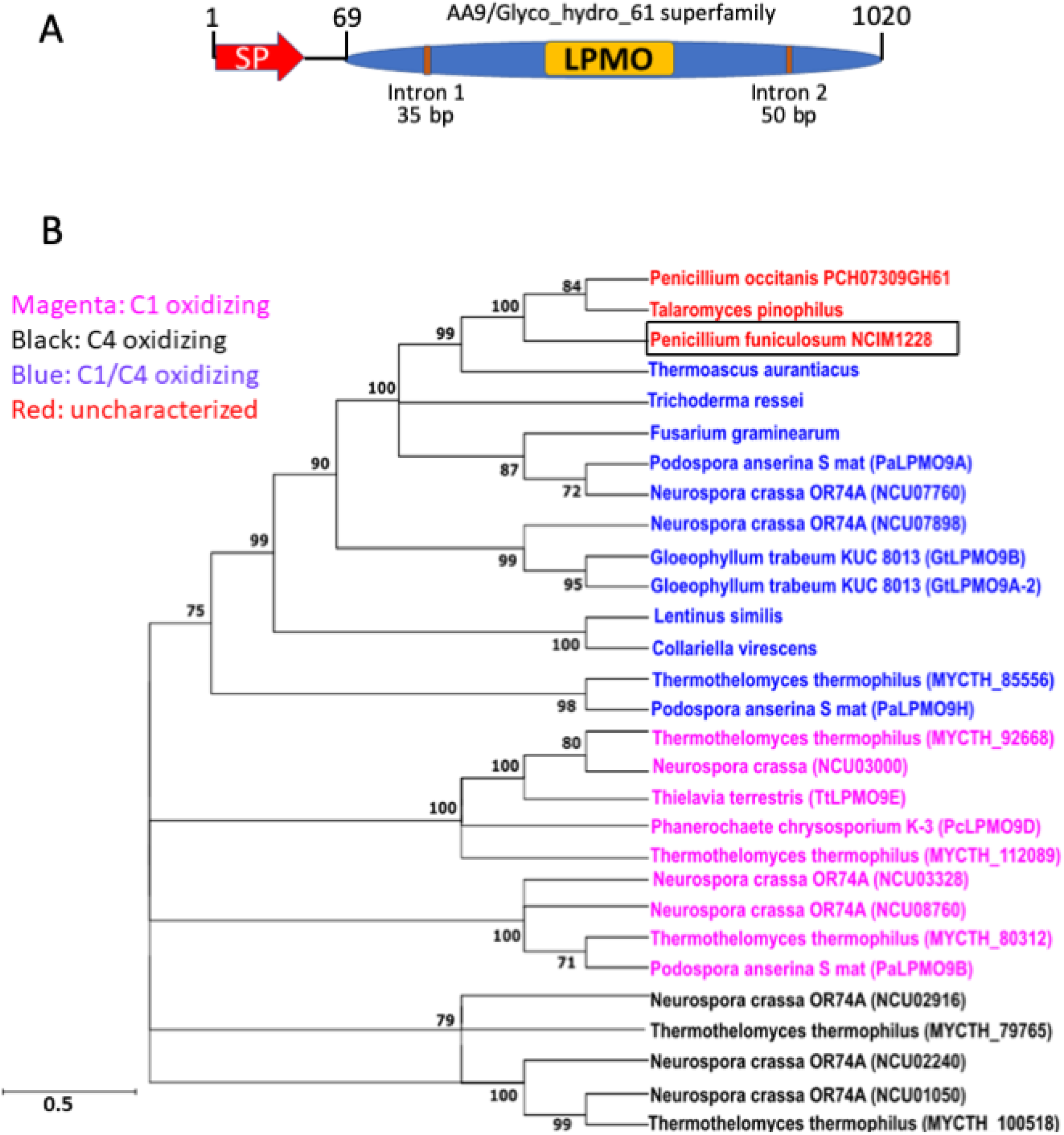
(A) Schematic representation of *P. funiculosum* LPMO and (B) phylogenetic tree of LPMO orthologs in fungi. Molecular Phylogenetic analysis was done by using the Maximum Likelihood method and the JTT matrix-based model. The tree with the highest log likelihood (−17992.93) is shown. Initial tree(s) for the heuristic search were obtained automatically by applying Neighbor-Join and BioNJ algorithms to a matrix of pairwise distances estimated using the Maximum Composite Likelihood (MCL) approach and then selecting the topology with superior log likelihood value. The tree is drawn to scale, with branch lengths measured in the number of substitutions per site. The bootstrap support corresponding to the numbers on the tree branches is calculated per 1000 bootstrap replicates. This analysis involved 29 nucleotide sequences. There was a total of 435 positions in the final dataset (71). Evolutionary analyses were conducted using MEGA X software.

**Fig. 2:**
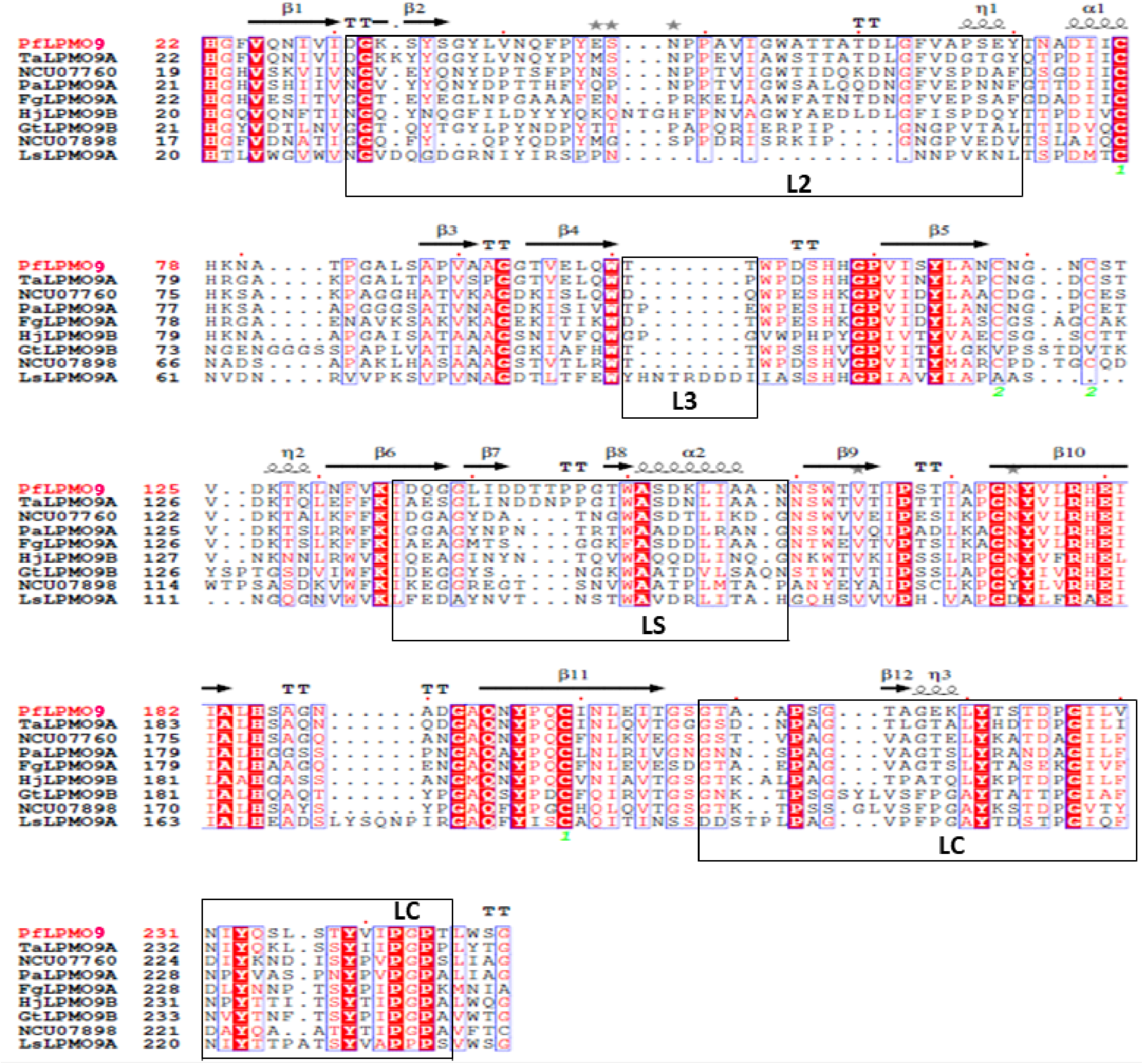
Multiple sequence alignment of PfLPMO9 with known C1/C4-oxidizing LPMO structures. The proteins included are as follows: NCU07760 (UniProtKB Q7S111), PaLPMO9A (UniprotKB B2B629), TaLPMO9A (UniprotKB G3XAP7), FgLPMO9A (UniprotKB I1REU9), HjLPMO9B (UniprotKB O14405), GtLPMO9B (UniprotKB Q7SA19)), and LsLPMO9A (Uniprot A0A0S2GKZ1). Fully conserved residues are shown in white on a red background. Blue frames indicate that more than 70% of the residues in the corresponding columns exhibit similar physicochemical properties (indicated as red residues on a white background). Black boxes indicate variable regions in the LPMO9 family with the corresponding names L2, L3, LC, and LS, which contribute to shaping the substrate-binding surface.

For detailed structural comparison, TaLPMO9A (PDB ID: 2YET), which showed closest structural similarity with PfLPMO9, and LsLPMO9A (PDB ID: 5ACI), one of the most characterized C1/C4 LPMO of the AA9 family whose crystal structure has been solved and made available in public repositories, were taken for consideration. To evaluate the amino acids involved in catalysis by this LPMOs, the modelled structures of PfLPMO9 and TaLPMO9 were docked with cellohexaose by taking the crystal structure of LsLPMO9A as the reference model (Fig. 3). The ligand-binding affinities of the LPMOs in terms of change in Gibbs free energy derived from Molecular Mechanics Generalized Born Surface Area (MMGBSA) calculations of the top conformations of PfLPMO9 and TaLPMO9A modelled complex are presented in Table 1 along with that of the crystal structure of LsLPMO9A. Surprisingly, we found a significantly lower change in Gibbs free energy for the different complexes of PfLPMO9 when compared with that of TaLPMO9A (Fig. S2). The lowest free energies obtained as calculated by the MMGBSA assay using Schrodinger suite (version Maestro 2019-4) were -46 kcal/mol and -31 kcal/mol for PfLPMO9 and TaLPMO9A, respectively (Table 1). This indicates that PfLPMO9 has a higher binding affinity of the enzyme for the substrate as compared to TaLPMO9A. However, LsLPMO9A was found to have a ΔG value closely similar to that of PfLPMO9, which was -49 kcal/mol (Table 1).

**Table 1:**
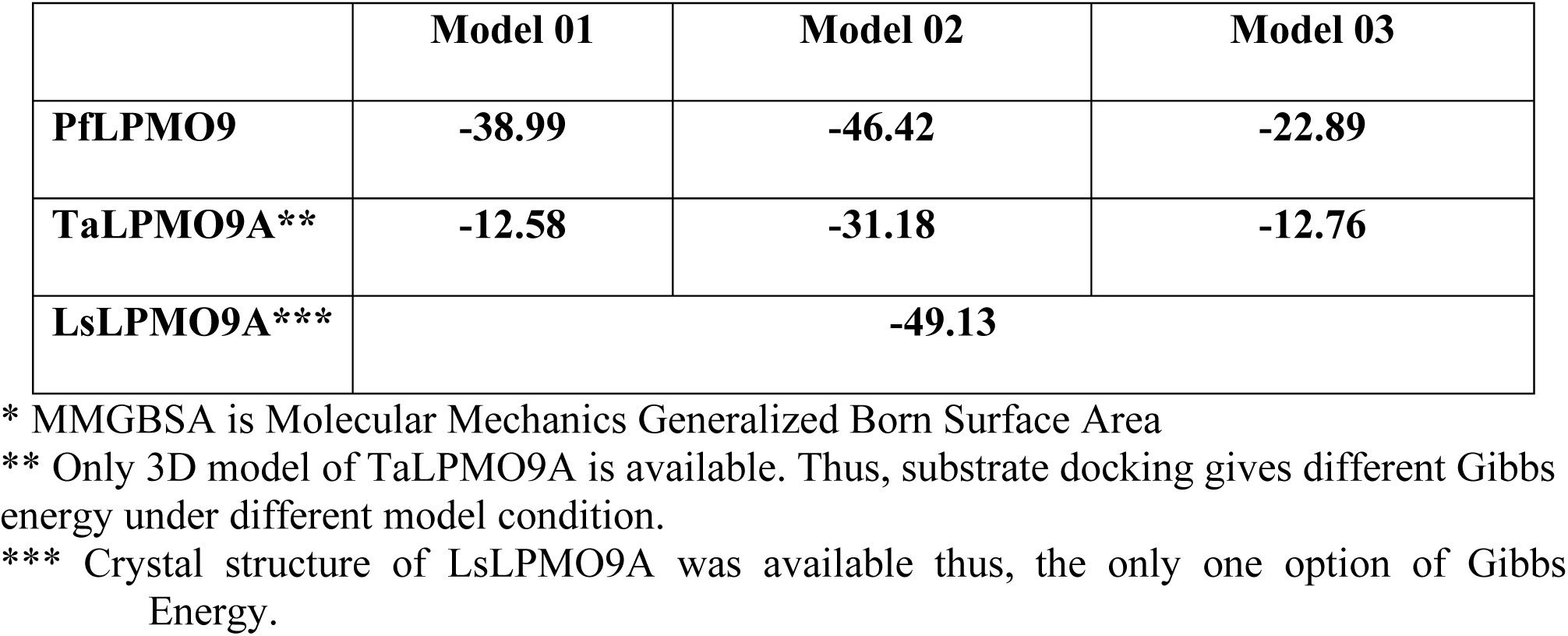
MMGBSA* ΔG binding energies (kcal/mol) of the different model complexes of PfLPMO9 and TaLPMO9A docked with the available crystal structure of LsLPMO9A-cellohexaose complex

**Fig. 3:**
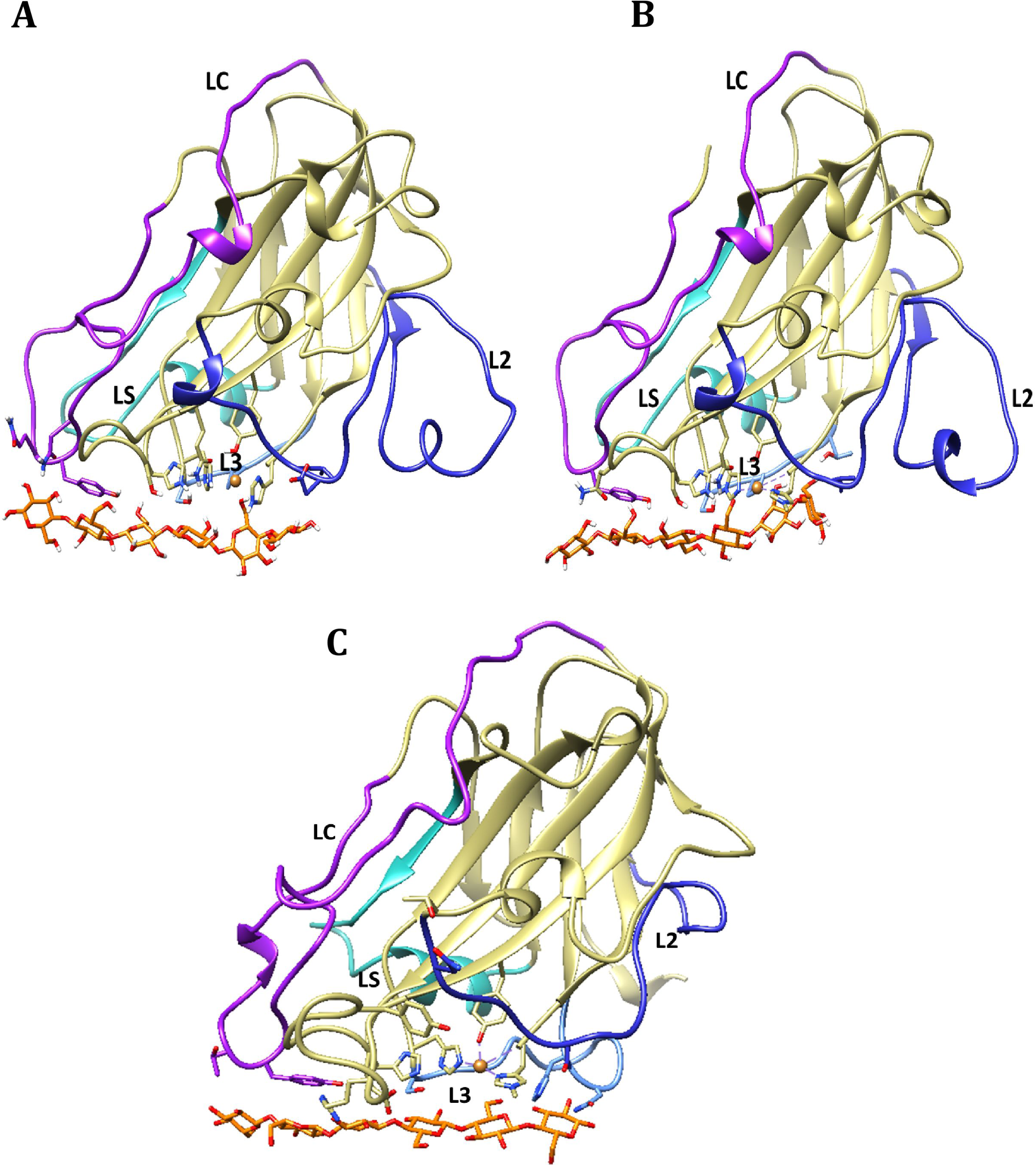
Comparative structures of (A) PfLPMO9, (B) TaLPMO9A docked with cellohexaose and (C) LsLPMO9A crystal structure in complex with cellohexaose. Variable regions in the LPMO9 family which contribute to shaping the substrate-binding surface are indicated with the corresponding names L2, L3, LC, and LS.

Furthermore, close observation of the mode of interaction of the amino acid residues at the active sites of the LPMOs via LIGPLOT (v.4.5.3) (35) revealed that PfLPMO9 had almost similar enzyme-ligand interaction with that of LsLPMO9A (Fig. 4). This might be the reason for the similar lower binding energies and molecular interactions with substrate obtained with both complexes as compared with that of TaLPMO9A (Table S1), even if PfLPMO9 and TaLPMO9A share more structural similarity. This may thus suggest that the nature and arrangement of the amino acid residues in the binding pocket of proteins are essential for their binding affinity. The superimposition images of the obtained PfLPMO9 structure with TaLPMO9A structure as well as with LsLPMO9A structure are shown in Fig. 5A and 5B, respectively. We found a very close similarity in the overall folds of PfLPMO9 and TaLPMO9A, while PfLPMO9 and LsLPMO9A showed a different folding pattern due to variations of the amino acids in both LPMOs as seen earlier from the multiple sequence alignment (Fig. 2). The structure models indicated two possible disulphide bonds in both PfLPMO9 (Cys118–Cys122 and Cys77–Cys199) and TaLPMO9A (Cys97–Cys101 and Cys56–Cys178), as shown in Fig. 6A, which might help in the stabilization of their tertiary structures. On the other hand, only one disulphide bond was reported for LsLPMO9A (Cys41-Cys167) (36), as indicated in Fig. 6B. Furthermore, we noticed only two critical changes in the amino acid residues when PfLPMO9 and TaLPMO9A were superimposed (Fig. 6A), while several changes were seen with the superimposition of PfLPMO9 with LsLPMO9A (Fig. 6B). The presence of threonine, a polar amino acid, at position 81 in PfLPMO9 in place of proline in TaLPMO9A was observed from the alignment. Also, glycine was seen in PfLPMO9 at position 167 in place of glutamine in TaLPMO9A. *Thermoascus aurantiacus* is a well-known fungus for the production of thermophilic lignocellulolytic enzymes (37, 38). More importantly, its LPMO (TaLPMO9A) has recently been reported to be highly thermostable where it was shown to retain 72% of its activity after 45 min of incubation at 100 °C, thereby holding great prospect in the enzymatic deconstruction of cellulosic biomass at elevated temperatures (39). It may thus be inferred that PfLPMO9 could be a thermotolerant enzyme based on its high structural similarity to TaLPMO9A in terms of the arrangement of the amino acid residues and overall folding.

**Fig. 4:**
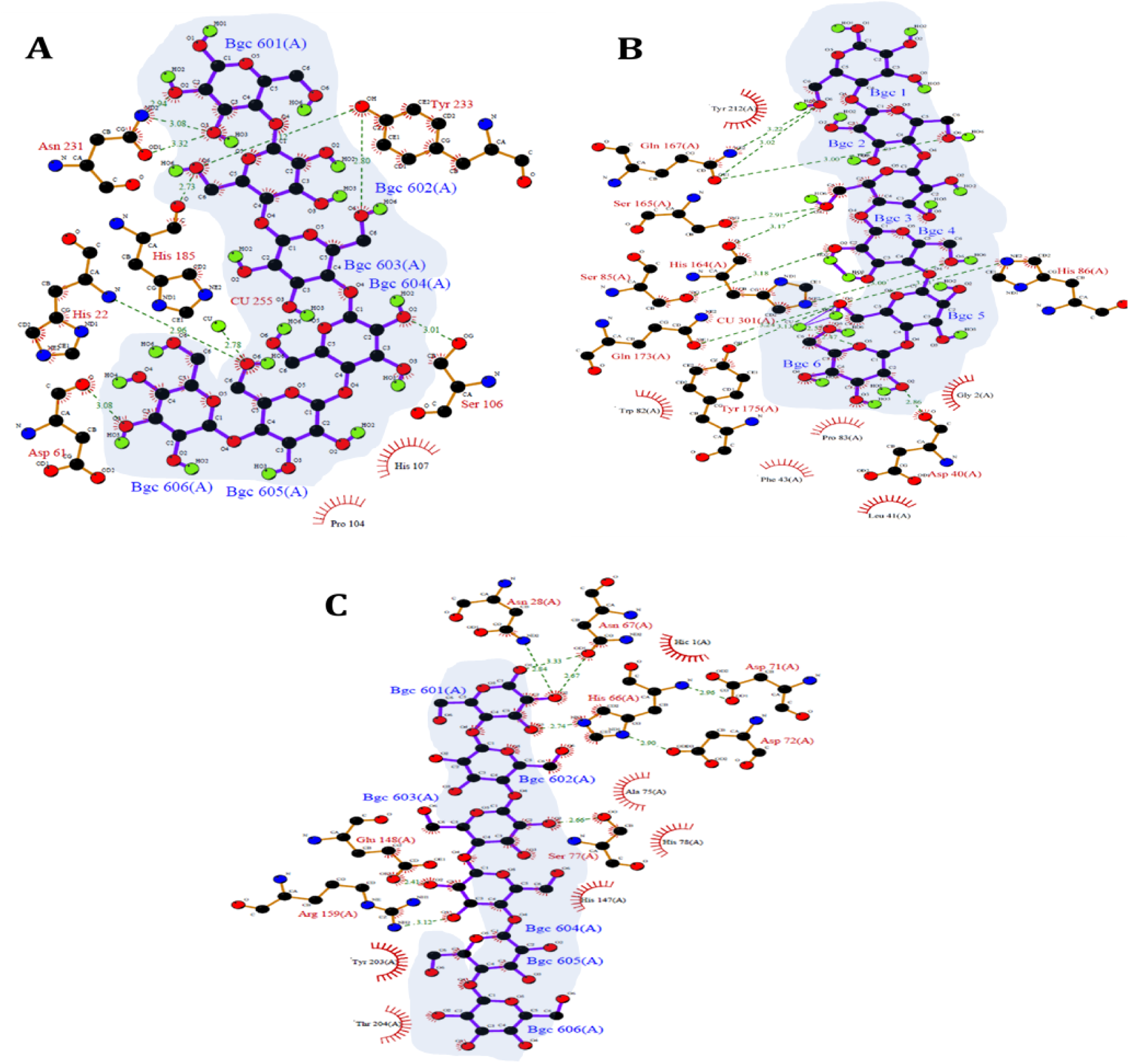
Ligplot analysis showing the molecular interactions and amino acid residues involved in substrate binding of (A) PfLPMO9, (B) TaLPMO9A and (C) LsLPMO9A with cellohexaose. (images are for complexes having lowest MMGBSA ΔG bind values). The arrangements of glucose rings in the cellohexaose ligand are indicated in blue background.

**Fig. 5:**
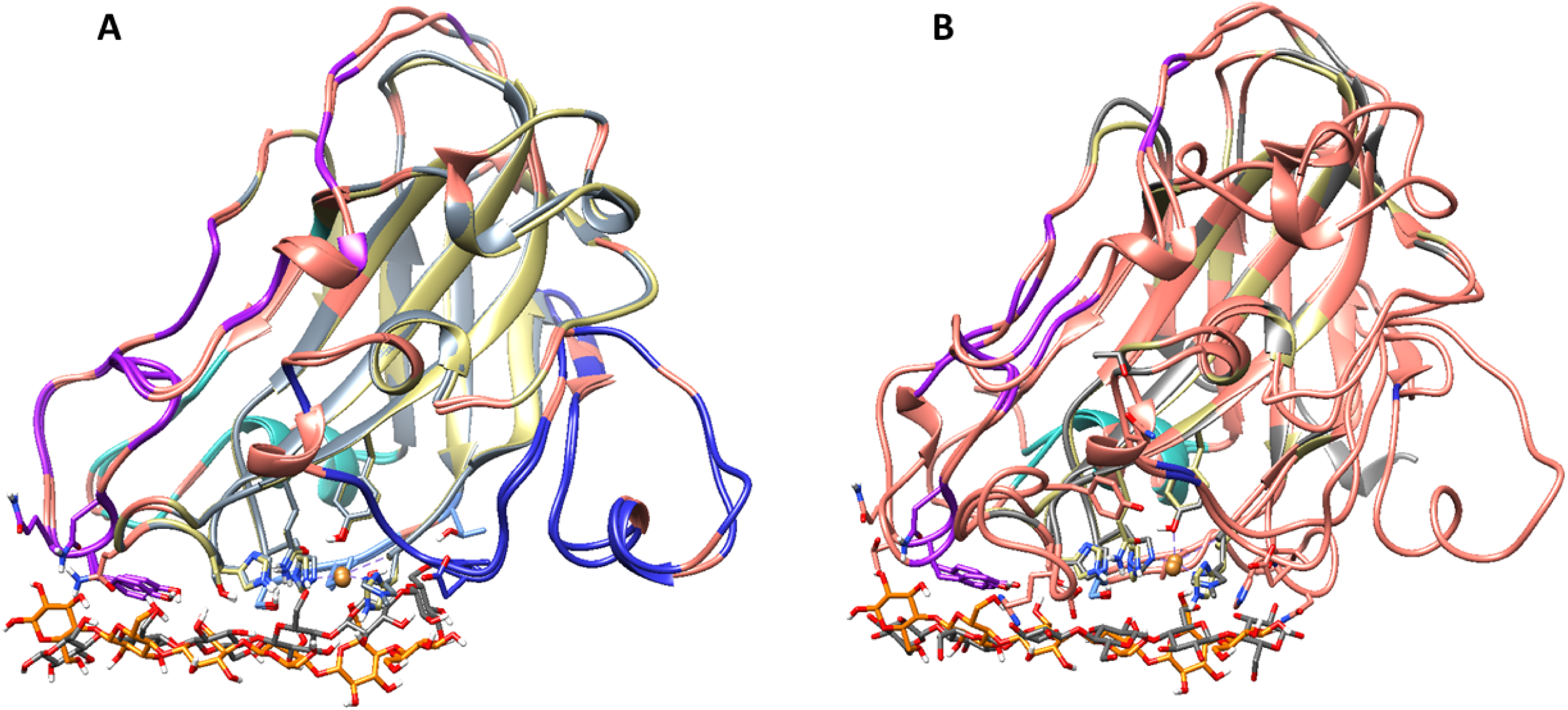
Conformational images for the superimposition of (A) PfLPMO9 with TaLPMO9A and (B) PfLPMO9 with LsLPMO9A in complex with cellohexaose. Non-conserved residues for both complexes showing regions which are non-identical are indicated in brown colour. Amino acid residues common to PfLPMO9 alone are highlighted in gold while amino acid residues for TaLPMO9A and LsLPMO9A only are shown in grey colour.

**Fig. 6:**
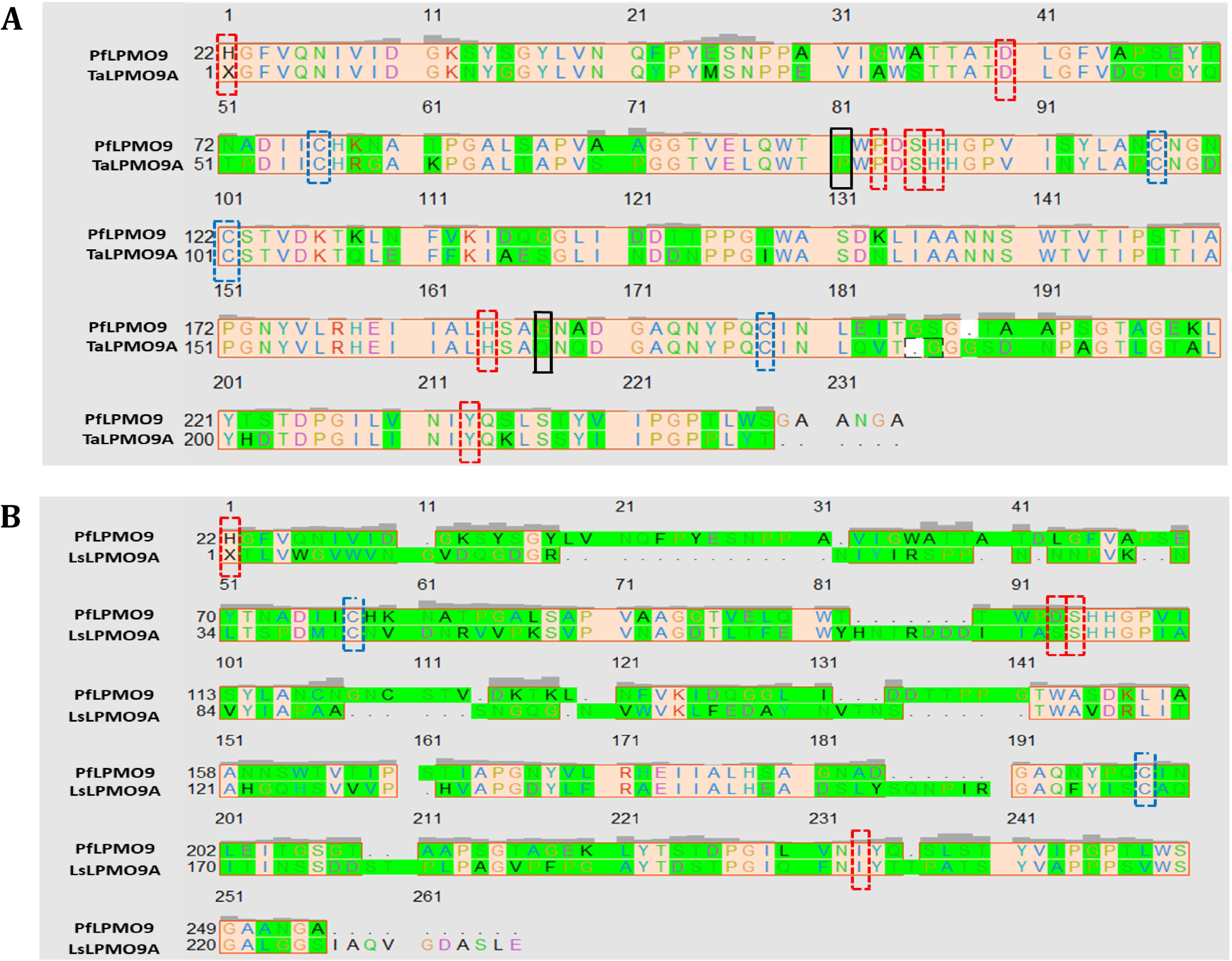
Comparative analysis of the amino acids involved in catalysis by (A) PfLPMO9 and TaLPMO9A; (B) PfLPMO9 and LsLPMO9A after superimposition of their 3-D structures. Similar amino acid residues present at the active sites of both enzymes are indicated in red-dashed boxes. The cysteine residues responsible for the disulphide bridges in the structures are indicated in blue-dashed boxes. Different amino acid residues belonging to the same group are shown in the green background, while the amino acids belonging to different groups are indicated in black dashed boxes.

To evaluate if PfLPMO9 is indeed a thermotolerant enzyme, we performed a thermostability study for the enzyme by incubating the enzyme without substrate at temperatures ranging from 50 to 80 °C for 300 min. Aliquots of the enzyme were withdrawn at 30 min intervals and used to determine their residual activity after incubation (Fig. 7). The thermostability profile of PfLPMO9 showed that the enzyme was highly thermostable as it retained above 50% of its original activity after 60 min of incubation at elevated temperatures up to 80 °C (Fig. 7). The enzyme showed stable activity at 50 and 60 °C where it was found to have a half-life of 87 and 17 h, respectively (Fig. 7, Table 2). The results showed that 96% of its original activity was retained at 50 °C after 300 min (5 h) of incubation while ∼40% residual activity was obtained at 70 °C after this time of incubation. At 80 °C also significant activity was observed with T_1/2_ of 1.1 h. Interestingly, we found this PfLPMO9 to have comparable thermostability property with that reported from *Scytalidium thermophilum* (PMO9D_SCYTH) and *Malbranchea cinnamomea* (PMO9D_MALCI) with half-life of 23 h at 60 °C (40). Similarly, it exhibited higher thermostability than that reported for different *Penicillium* strains as well as some other fungal species such as *Trichoderma reesei, Pestalotiopsis sp and Aspergillus fumigatus* whose half-lives were in the range of 8 – 13 h at 50 oC (41–43). Utilization of thermostable enzymes, including LPMOs, are of interest industrially as saccharification performed at high temperature decreases the risk of microbial contamination and enables enhanced saccharification rates at higher substrate loading due to reduced slurry viscosity (44). The ability of PfLPMO9 to retain significant enzymatic activity at higher temperatures validates its thermotolerance characteristics as predicted from our *in-silico* analysis. A combination of impressive thermostability and higher binding affinity features of PfLPMO9 deciphered in this study suggest it as a promising enzyme for utilization in biomass saccharifications.

**Table 2:** Half-life of PfLPMO9 after incubation at different temperatures.

|  | 50 °C | 60 °C | 70 °C | 80 °C |
| --- | --- | --- | --- | --- |
| $T_{1/2}$ (min) | 5266.32 | 1043.86 | 230.53 | 62.80 |
| $T_{1/2}$ (h) | 87.77 | 17.40 | 3.84 | 1.05 |

**Fig. 7:**
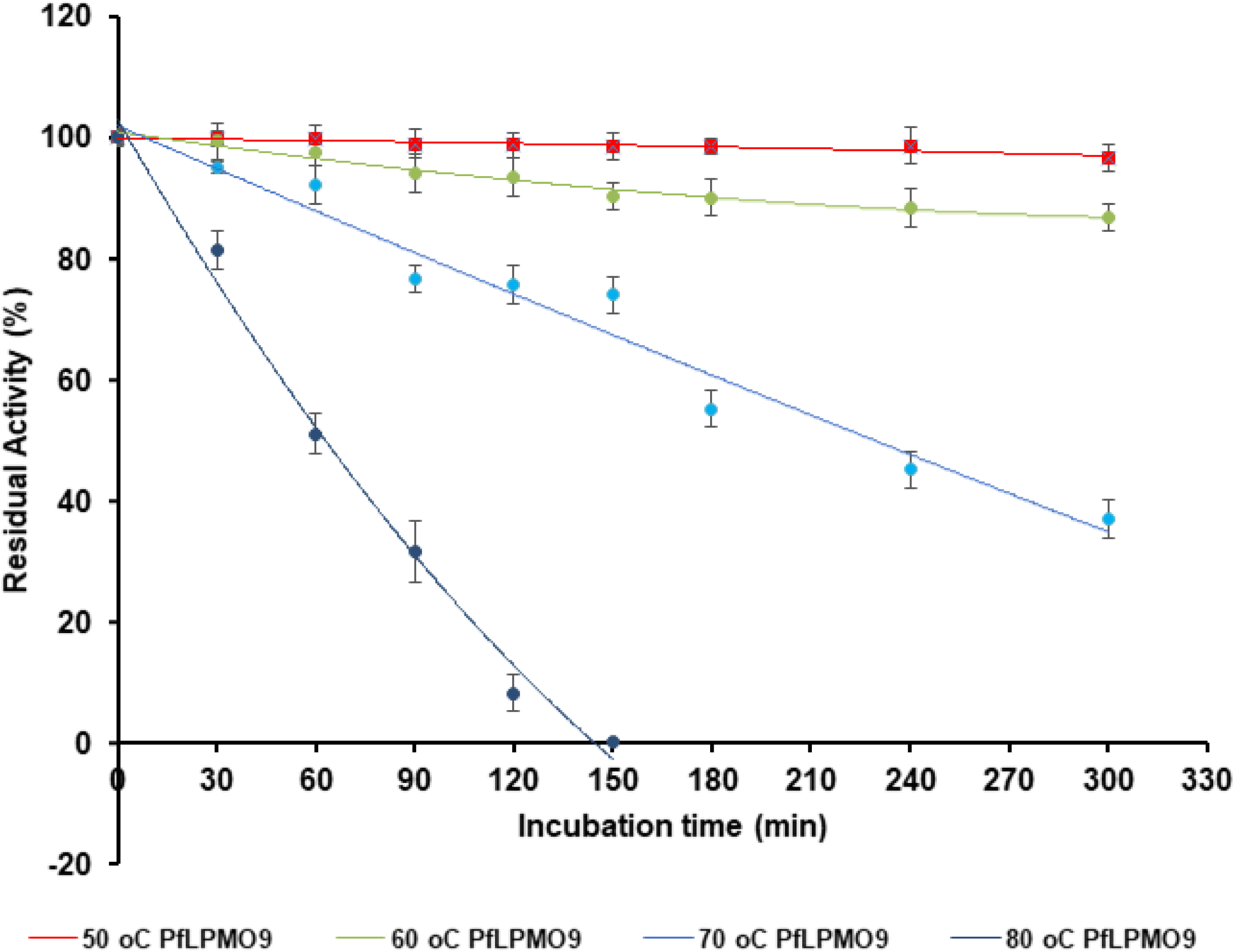
Thermostability profile of LPMO enzyme from *P. funiculosum*. The residual activities were calculated using the maximum enzyme activity as 100%. The half-life of the enzyme at each temperature was calculated from the slope of the curves (Error bars represent mean ± standard deviation of triplicate determination).

### Regulation of PfLPMO9 gene expression in *Penicillium funiculosum*

To investigate the functional role of PfLPMO9 in the degradation of cellulosic biomass, the fungus was grown on cellulosic and hemi-cellulosic biomasses and its transcriptional regulation was evaluated by comparing its RNA level with that grown on glucose as carbon source. Transcriptional analysis of PfLPMO9 gene was performed using RNAseq data available for this fungus grown on various carbon source in our lab. The LPMO was found to be upregulated when the fungus was cultivated on crystalline cellulose (Avicel) and hemicellulose (wheat bran) as well as on alkali-pretreated wheat straw as compared to that when grown on glucose, a readily metabolizable carbon source (Table 3). It was very striking to see that the LPMO was most upregulated in the presence of Avicel with log2 fold change of 7.73 with respect to glucose, strongly suggesting its possible unique role in the degradation of crystalline cellulose as compared to other biomasses tested (Table 3). The extent of upregulation of PfLPMO9 gene in response to crystalline cellulose also closely matched with that of the CBH1 gene (Table 3), whose product Cellobiohydrolase I is well known to act upon crystalline cellulose. The results from the transcriptomic analysis indicate that improving the abundance of PfLPMO9 in the secretome may positively facilitate an improvement in its saccharification efficiency on very recalcitrant and crystalline biomass as earlier suggested from our previous proteomic studies (19, 28).

**Table 3:**
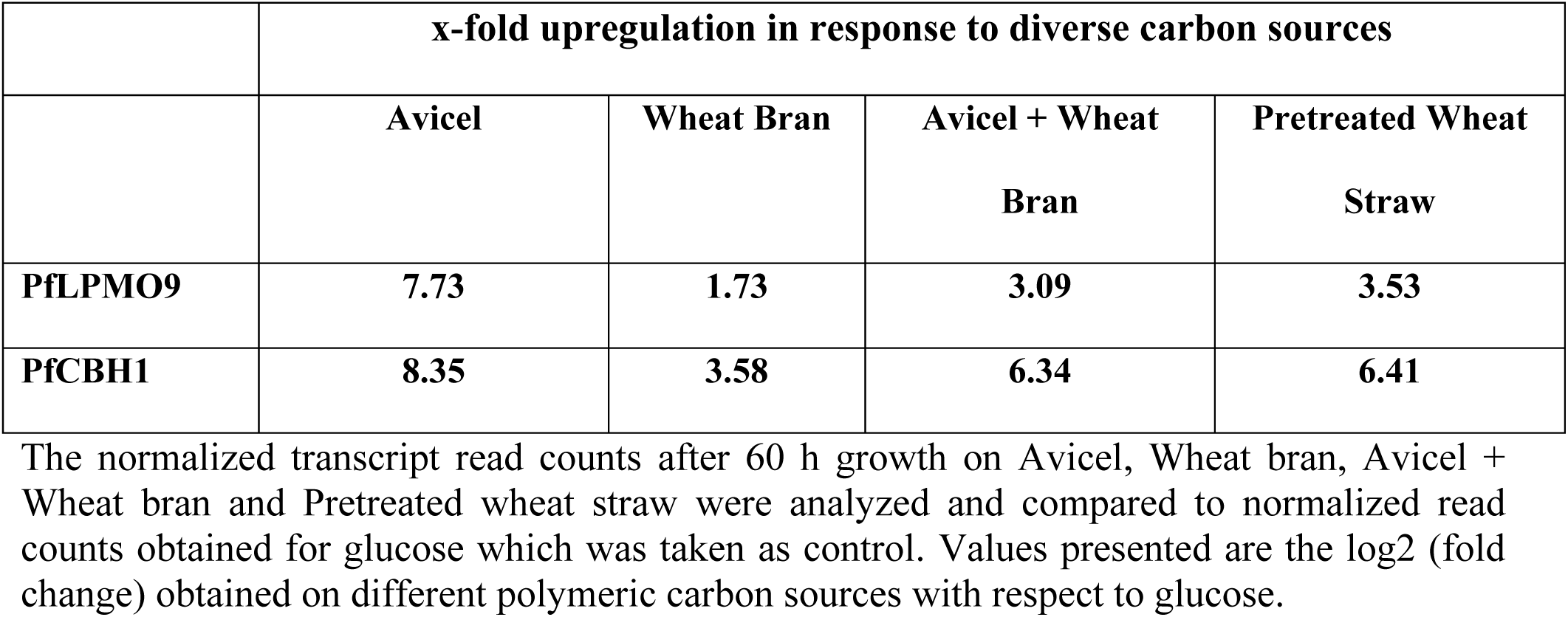
Transcriptional regulation of genes coding for PfLPMO9 and PfCBH1 in response to diverse carbon sources

### Recombinant overexpression of PfLPMO9 in PfMig1^88^ strain to improve LPMO activity

We decided to insert additional copy of PfLPMO9 gene in the background of Mig1-repressor deleted strain of *P. funiculosum*, i.e., *Pf*Mig1^88^, as it was shown to produce higher quantity of cellulolytic enzymes in our earlier study (25). To overexpress the PfLPMO9 gene in *Pf*Mig1^88^ strain, the expression vector containing the endogenous gene along with its promoter and terminator was constructed using the backbone of pBIF binary vector. The 3.0 kb region containing LPMO9 gene (Fig. 8A) was amplified from the genome of the parent NCIM1228 strain and cloned into the pBIF vector to generate the pOAO1 plasmid. The pOAO1 recombinant plasmid was confirmed by restriction digestion before transformation into *Pf*Mig1^88^ strain (Fig. S3A) using the AMTM procedure. The hygromycin-resistant transformants obtained were then subsequently analyzed (Fig. 8B). The transformants were first confirmed by PCR (Fig. S3B), and Southern blot was then carried out for five of PCR-positive transformants to further check the integration and copy number. For this, genomic DNA from each strain was digested with *Xho*I and *Hind*III and probed with a 621-bp LPMO gene fragment (Fig. 8C). Results showed that all the transformants possessed more than one bands, in contrast with the control (*Pf*Mig1^88^) that had only one single band, further verifying the insertion of an additional copy of LPMO cassette in the genome of *Pf*Mig1^88^.

**Fig. 8:**
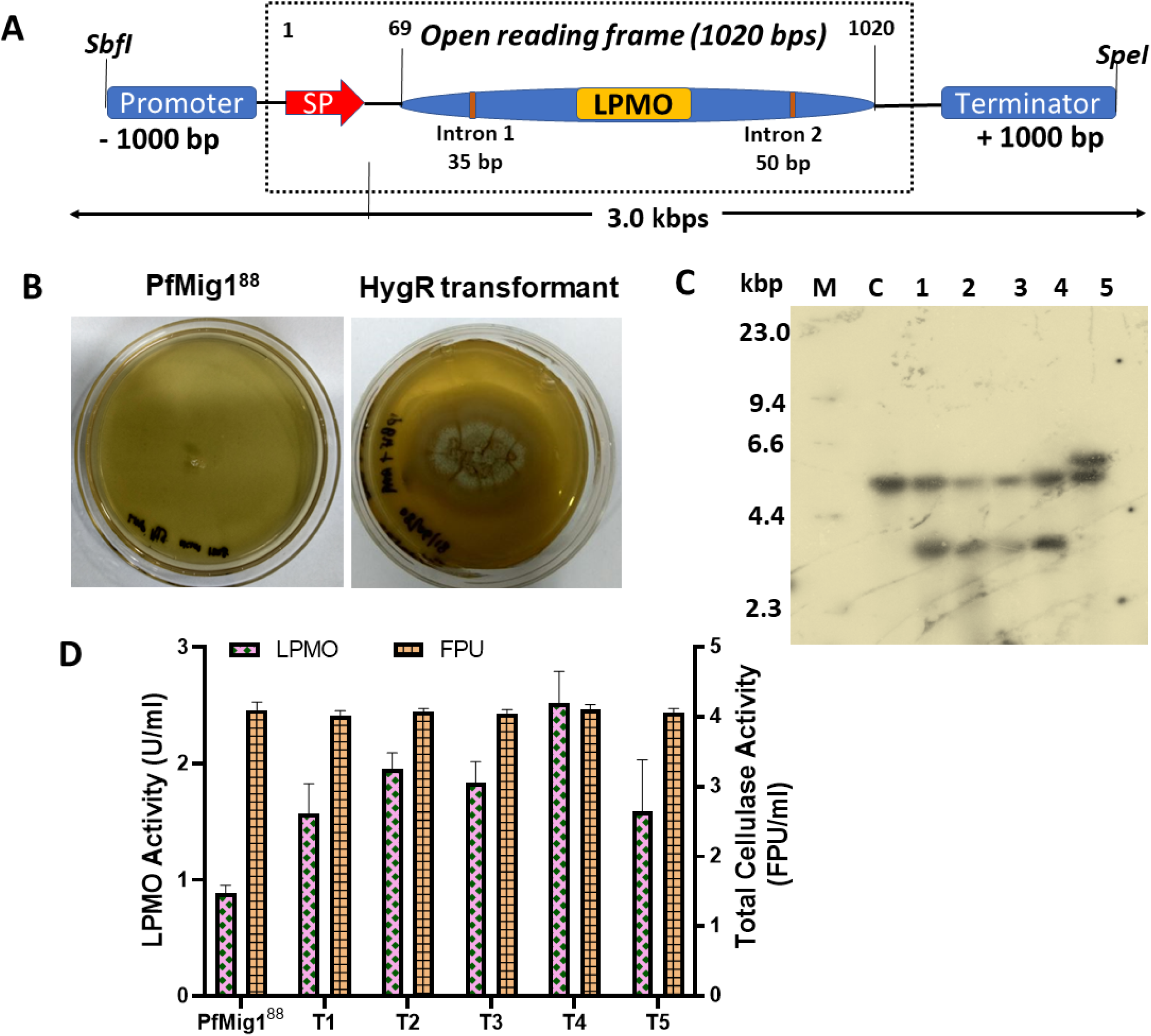
Construction of LPMO expression cassette and its overexpression in *Pf*Mig1^88^. (A) Schematic diagram showing the assembly of LPMO cassette from *P. funiculosum* NCIM1228 genome. (B) Transformants of pOAO1 after AMTM transformation in *Pf*Mig1^88^. Transformants were selected on 100 µg/ml hygromycin. (C) Southern blot of transformants confirmed by PCR. M is *Hind*III lambda DNA size marker used; Lane C indicates genomic DNA of non-transformed parental strain; Lanes 1-5 represent genomic DNA from transformants with LPMO cassette integration; (D) The LPMO and FPase activities in the fermentation broth of *Pf*Mig1^88^ and five transformants.

In order to evaluate the expression level of LPMO gene in the LPMO-overexpressing transformants, the five transformants confirmed by Southern hybridization were cultivated in cellulase inducing media for five days. The LPMO activity of the transformants was measured using the culture supernatant and was compared to the parent strain, *Pf*Mig1^88^ (Fig. 8D). The results showed a significant increase in LPMO activity in the range of 155 – 203% in all the transformants when compared to the parent strain. The transformant T4 showed the maximum activity of 2.68 U/ml which was more than 200% increase over that of *Pf*Mig1^88^ strain (0.88 U/ml). Variation in enzyme activity across the transformants as determined by the Amplex red assay might be due to the difference in integration loci of the expression cassette in *Pf*Mig1^88^ genome as earlier reported (45, 46). However, when the total cellulase activity of the transformants was evaluated using the filter paper assay to see if there would be any corresponding impact on the overall cellulase activity, no significant difference in the activity of the transformants was observed as compared to that of the parent strain (Fig. 8D). The indifference could be due to the absence of reducing agents which serves as electron donors required by the LPMO enzyme to act on pure cellulose in the filter paper assay mixture (15). Nevertheless, the remarkable increase in LPMO activity recorded in all the transformants over that of the parent strain indicates that the gene was successfully overexpressed and the LPMO amount in the *Pf*Mig1^88^ cellulase system was significantly improved.

### Simultaneous overexpression of PfLPMO9 and PfCBH1 in *P. funiculosum* Mig1^88^ strain to improve cellulase system in its secretome

Although LPMO activity got significantly improved with the addition of an extra copy of LPMO gene, more effort needed to be put in place to improve the quality of cellulase system in the secretome in order to achieve higher degradation of cellulosic biomass. Since CBH1 happens to be one of the vital hydrolytic enzymes of particular importance participating in crystalline cellulose deconstruction, the impact of overexpression of the CBH1 on the quality of total cellulase system of the LPMO-transformant was next examined.

In addition to the construction of the dual transformant, a fungal strain was also engineered where only CBH1 gene along with its promoter and terminator was integrated in the genome of *Pf*Mig1^88^ strain to assess the impact of CBH1 overexpression alone on the cellulase system (Supplementary file S1; Fig. S4A-E). About 57 - 62% increase in Avicelase activity was found across the five transformants over that of *Pf*Mig1^88^. The results also showed up to 25% increase in the filter paper unit of the transformants, where the maximum FPase achieved was5.1 FPU/ml for CBH1-transformant T3 (Fig. S4F)

To engineer the fungal strain for overexpression of both LPMO and CBH1, a systematic approach was utilized for constructing the desired pOAO5 vector containing the LPMO/CBH1 expression cassette (Fig. 9A), as described in Materials and Methods. For fungal transformation, the pOAO5 plasmid was confirmed by restriction digestion before transformation into *Pf*Mig1^88^ strain (Fig. S3C). Hygromycin-resistant transformants obtained were analyzed by PCR to confirm the integration of LPMO/CBH1 cassette into the genome (Fig. 9B, Fig. S3D). Similarly, to further confirm the integration and copy number, Southern blot was carried out for all selected transformants (Fig. 9C). Results showed that all the transformants possessed more than one band in contrast with the control (*Pf*Mig1^88^) strain which had a single prominent band, further verifying the insertion of an additional copy of the LPMO/CBH1 cassette in the genome of *Pf*Mig1^88^. Transformant T5 showed two faint bands at similar locations which may be due to experimental error leading to lower loading of digested genomic DNA. Four out of the five transformants had only one additional band, suggesting a single-copy insertion of LPMO/CBH1 cassette in the genome, while one of the transformants (T1) had two copies of the integrated cassette (Fig. 9C).

**Fig. 9:**
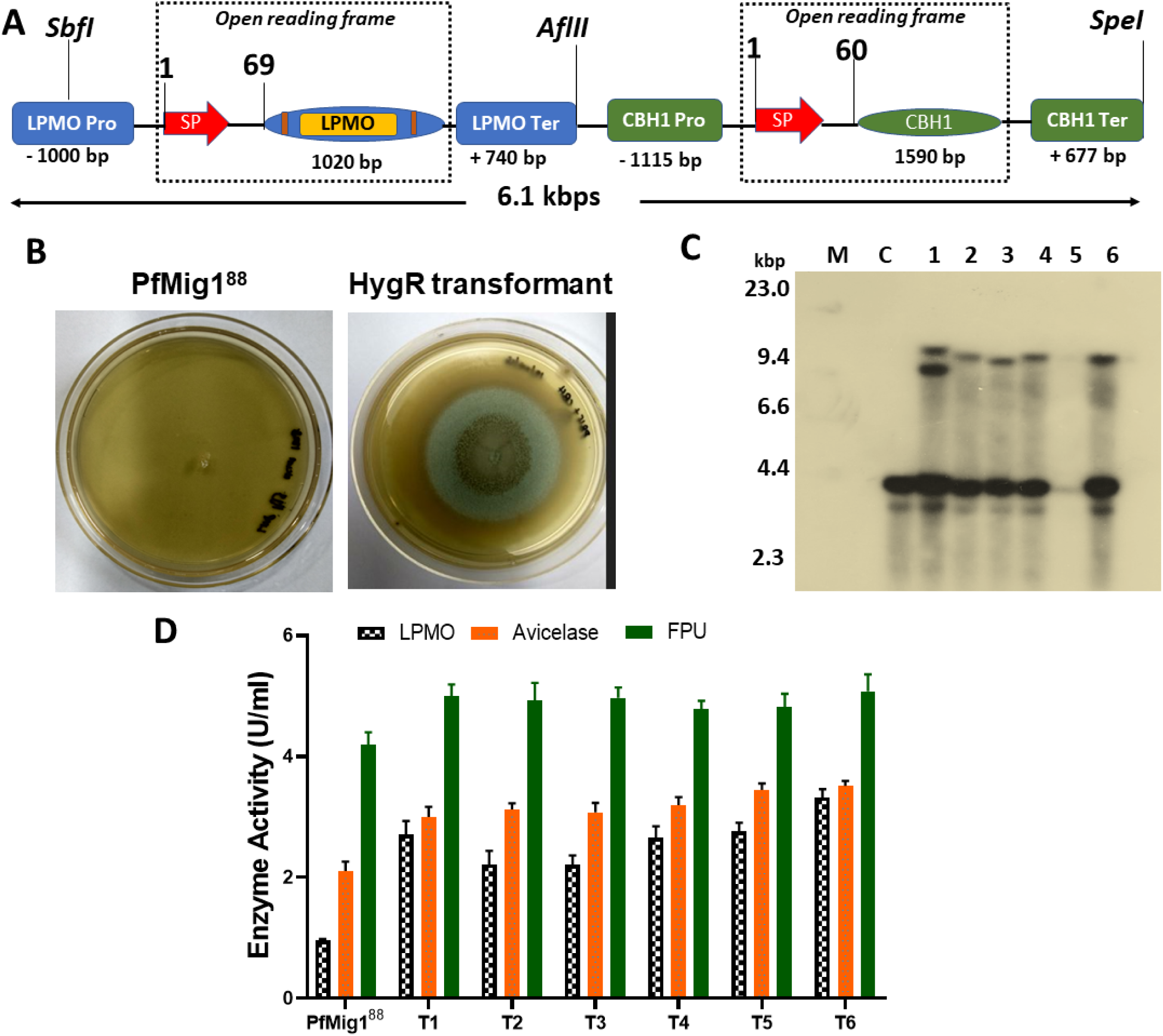
Simultaneous overexpression of LPMO and CBH1 in *Pf*Mig1^88^. (A) Schematic diagram showing the construction cassette for dual overexpression of LPMO and CBH1 genes from *P. funiculosum* NCIM1228. (B) Transformants of pOAO5 after AMTM transformation in *Pf*Mig1^88^. Transformants were selected on 100 µg/ml hygromycin. (C) Southern blot of transformants confirmed by PCR. M is HindIII lambda DNA size marker used; Lane C indicates genomic DNA of non-transformed parental strain; Lanes 1-5 represent genomic DNA from transformants with LPMO/CBH1 cassette integration; (D) The enzymatic profile of the overexpressed enzymes in the fermentation broth of *Pf*Mig1^88^ and six transformants.

To further assess the effect of this simultaneous overexpression on the expression level of cellulolytic enzymes by the transformants, the six transformants confirmed positive by PCR for the integration of the LPMO/CBH1 expression cassette were cultivated in cellulase inducing media for five days. The resulting secretome from the strains was recovered and used for enzyme assays. The LPMO, cellobiohydrolase and total cellulase activities of the transformants were measured and compared with that of parent *Pf*Mig1^88^ strain. As expected, based on the results from the specific overexpression of the enzymes, all the transformants screened demonstrated an enhancement in all the activities when compared with the parent strain as presented in Fig. 9D. The results showed that there was about 130 – 212% increment in LPMO activity over that of *Pf*Mig1^88^ across the transformants. Similarly, we found about 40 – 66% increase in Avicelase activity and 14 - 20% increase in filter paper unit when compared with *Pf*Mig1^88^ (Fig. 9D). Although there was variation in enzyme activities tested across all the transformants, the highest production of all the enzymes was found with transformants T6 in which activities of LPMO, Avicelase and FPase were 3.3 U/ml, 3.51 U/ml and 5.2 U/ml respectively (Fig. 9D). It was surprising to see that there was no difference between the enzyme activities of the transformants with single-copy integration and that of transformant T1 which had two additional copies of the LPMO/CBH1 cassette. This suggested that the integration locus other than the gene copy numbers might have also affected the expression of the cellulolytic enzymes (47).

### Comparative transcriptional and translational analysis of cellulolytic enzymes in NCIM1228, *Pf*Mig1^88^ and the engineered strains

Conventionally, the transcriptional regulation of protein synthesis is key to understanding the mechanism of enzyme secretion by microorganisms. Since all the mutants generated in this study were earlier verified by PCR using specific primers for the integration of the overexpressed enzymes in the genome, the transcript abundance of each overexpressed genes at the mRNA level also needs to be examined to gain more understanding on the overall regulation of the cellulase system. Furthermore, to understand whether the overexpression of the LPMO and/or CBH1 had any impact on the expression of other key cellulolytic enzymes, we also checked the transcript level of some of the major cellulases/hemicellulases known to be produced by NCIM1228 under a de-repressing condition (Table 4) (28). For this, cultures of NCIM1228, *Pf*Mig1^88^ and the best performing transformants of *Pf*OAO1 (overexpressing LPMO in the background of *Pf*Mig1^88^), *Pf*OAO2 (overexpressing CBH1 in the background of *Pf*Mig1^88^) and *Pf*OAO3 (overexpressing LPMO as well as CBH1 in the background of *Pf*Mig1^88^) were grown in 4% Avicel for 48 h to get good mycelia growth. Transcript levels of all the five strains under de-repressing conditions were determined by real-time PCR with tubulin as control, and the relative fold change was normalized to NCIM1228 since it was the original strain from which all other mutants were generated. For LPMO overexpression, we observed a 5-fold increase in LPMO transcript for *Pf*Mig1^88^ while there was 18 and 26-fold increase in transcripts for *Pf*OAO1 and *Pf*OAO3 strains. As expected, there was no difference in the LPMO transcript level for *Pf*OAO2 transformant when compared to that of *Pf*Mig1^88^ (Fig. 10A). Likewise, for CBH1 overexpressing strains, about 4, 30 and 39-fold increase in CBH1 transcript was seen for *Pf*Mig1^88^, *Pf*OAO2 and *Pf*OAO3 strains, respectively, while there was no change in the CBH1 transcripts of *Pf*OAO1 (Fig. 10A). When the transcript level for β-glucosidase (BGL) gene in all the strains was checked, the results showed a 5-fold increase in BGL expression for *Pf*Mig1^88^ and *Pf*OAO1, while there were 10 and 11-fold increase recorded for *Pf*OAO2 and *Pf*OAO3 mutants, respectively. The increase in the expression level of BGL in the two CBH1 transformants could be as a result of the increase in cellobiohydrolase level in the transformants. The increased cellobiohydrolase level in the media may yield more cellobiose, thereby providing a signal to the cells to produce more BGL required to hydrolyze this cellobiose into glucose. However, no significant difference between the parent *Pf*Mig1^88^ strain and all the mutants was observed in the expression level of endoglucanase (EG) and xylanase (XYL) (Fig. 10A).

**Table 4:**
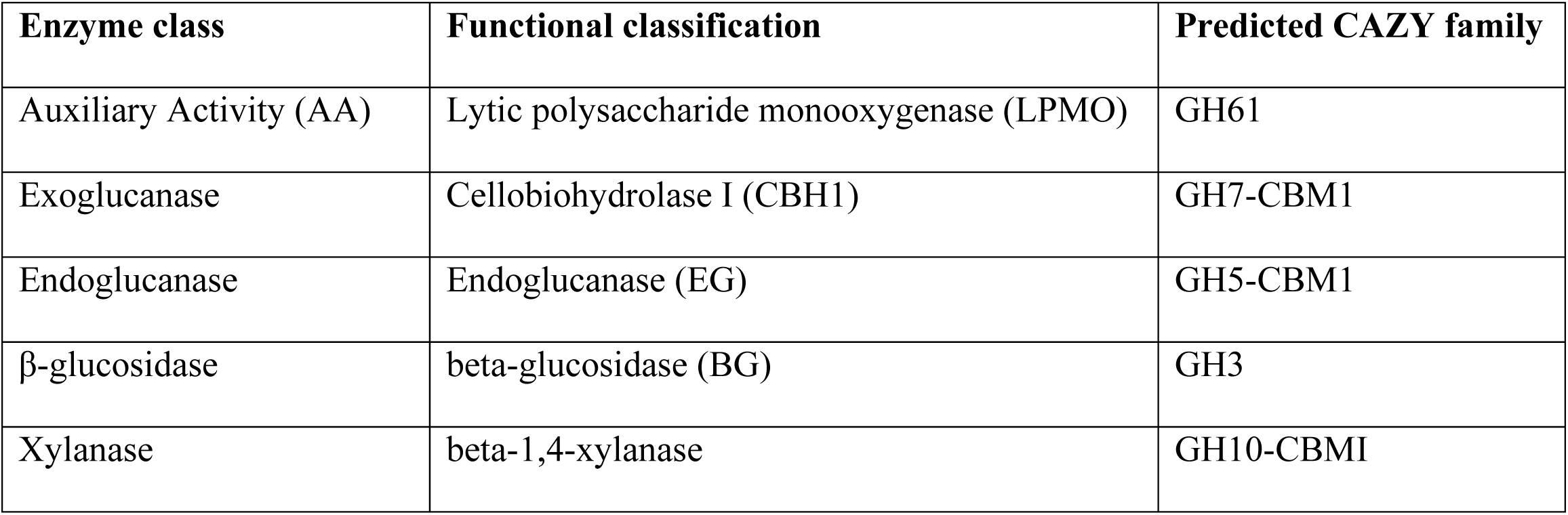
List of CAZymes whose transcript levels were monitored in *P. funiculosum* NCIM1228, PfMig1^88^ and the resulting transformants

| Enzyme class | Functional classification | Predicted CAZY family |
| --- | --- | --- |
| Auxiliary Activity (AA) | Lytic polysaccharide monooxygenase (LPMO) | GH61 |
| Exoglucanase | Cellobiohydrolase I (CBH1) | GH7-CBM1 |
| Endoglucanase | Endoglucanase (EG) | GH5-CBM1 |
| β-glucosidase | beta-glucosidase (BG) | GH3 |
| Xylanase | beta-1,4-xylanase | GH10-CBMI |

**Fig. 10:**
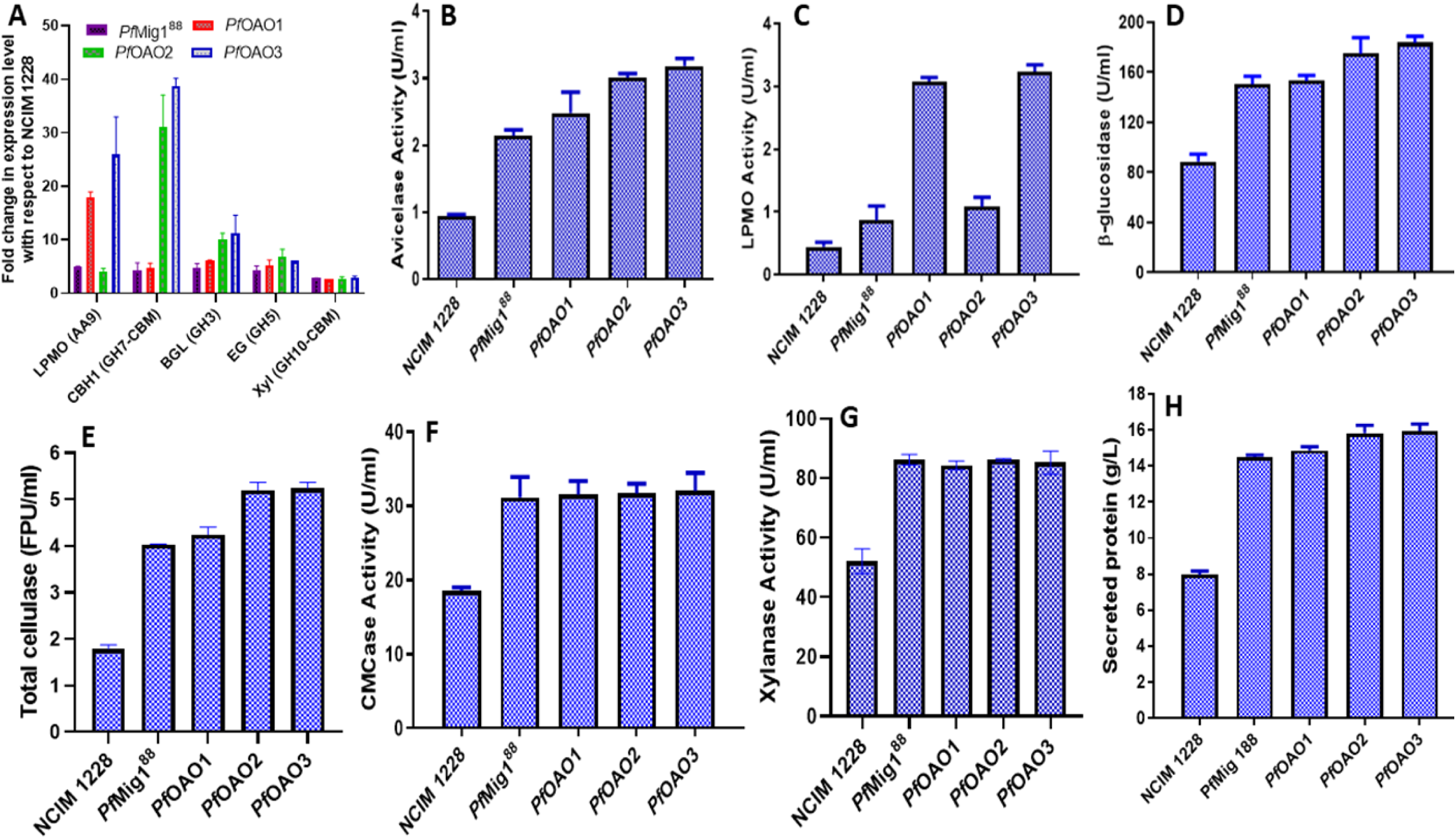
Determination of cellulase expression and activities in *P. funiculosum* NCIM1228, *Pf*Mig1^88^ and all the engineered strains. (A) The transcriptional expression of LPMO, cellobiohydrolase, endoglucanase, β-glucosidase, and xylanase in NCIM1228, *Pf*Mig1^88^ and all engineered strains were measured by quantitative real-time PCR after growing for 48 h in the presence of 4% Avicel. The expression levels were normalized to NCIM1228 and plotted. (B) The CBH1 activity determined using Avicel as the substrate; (C) LPMO activity determined using Amplex red; (D) β-glucosidase activity using pNPG; (E) The overall cellulase activity on filter paper; (F) The endoglucanase activity determined using CMC as the substrate; (G) The xylanase activity measured using beechwood xylan as the substrate; (H) Total secreted proteins in all strains. Data are represented as the mean of three independent experiments and error bars express the standard deviations.

To examine and compare the influence of LPMO and CBH1 overexpression on the overall cellulase system, cultures of NCIM1228, *Pf*Mig1^88^ and the best performing transformants of all engineered strains were grown in cellulase-inducing medium (CIM). The resulting supernatants containing the secreted enzymes were recovered from the mycelia and used for all enzyme assays. Individual cellulase activities of LPMO, CBH, EG, BGL, XYL and FPU for all strains were assayed, compared and presented in Fig. 10B to G. In accordance with the mRNA levels and earlier report (25), *Pf*Mig1^88^ strain showed a 2-fold increase in all enzyme activities evaluated, including the total extracellular protein. It was found that *Pf*OAO2 and *Pf*OAO3 showed ∼65% increase and 27% increase in Avicelase (Fig. 10B) and FPase activity (Fig. 10E), respectively, as compared to *Pf*Mig1^88^, while *Pf*OAO1 showed non-significant change. On the other hand, *Pf*OAO1 and *Pf*OAO3 showed 200% increase in LPMO activity, while *Pf*AOA2 did not show any significant change (Fig. 10C). In addition to increase in Avicelase activity, *Pf*OAO2 and *Pf*OAO3 also showed ∼18% increase in BGL activity and ∼ 9% increase in secreted protein concentration when compared with *Pf*Mig1^88^ (Fig. 10D, 12H), as indicated by transcriptome data. Furthermore, the increase in BGL production by *Pf*OAO2 and *Pf*OAO3 transformants relative to *Pf*Mig1^88^ was further confirmed by zymography analysis (Fig. S5). Not much change in CMCase activity and xylanase activity was observed in the transformants as compared to *Pf*Mig1^88^ (Fig. 10F-12G).

In addition, we enquired whether the specific activities of the overexpressed enzymes also increased besides volumetric activities. For this, the total protein in the secretome of each strain was determined and used to calculate their specific activities Table S2. The result showed a significant increase in the specific activity of LPMO in *Pf*OAO1 and *Pf*OAO3 strains relative to that of *Pf*Mig1^88^. Similarly, there was a significant increase in specific activities of CBH1, BGL and FPU in *Pf*OAO2 and *Pf*OAO3 overexpressing strains. However, there was no significant difference in the specific activities of EG and XYL across all the strains. The increase in specific activities of the key hydrolytic and oxidative enzymes seen in this study indicates that the performance of the enzyme mixture produced by the overexpressed strains have been enhanced when compared with *Pf*Mig1^88^ and may help in reducing the enzyme load required for biomass saccharification.

### Hydrolysis efficiency of cellulase enzyme complex with LPMO and CBH1 overexpression against acid-treated biomass at a high substrate loading

To assess the relevant industrial application of overexpression of these two key enzymes, the secretome from all the engineered strains as well as the parent strains was used to investigate their saccharification performance on acid pretreated wheat straw (PWS). The saccharification reaction was set up at 20% substrate loading of PWS using secretome of NCIM1228, *Pf*Mig1^88^, *Pf*OAO1, *Pf*OAO2 and *Pf*OAO3 at same enzyme loading of 30 mg/g dry biomass weight (DBW), and incubated at 50 °C for 96 h. Samples were collected every 24 h and analyzed for production of monomeric sugars (Fig. 11A and 11B). From the results, a linear increase in the concentration of monomeric sugars and hydrolysis efficiency was seen in all the strains with increasing time until the 72 h after which no appreciable increase in sugar concentration was observed. At 72 h time point, about 27% holocellulose (cellulose + hemicellulose) conversion was obtained in case of NCIM1228 secretome whereas more than 64% of the pretreated biomass was hydrolyzed by *Pf*Mig1^88^ secretome (Fig 11A); and the total monomeric sugars (glucose + xylose) recorded were 37.5 g/L and 90 g/L, respectively (Fig. 11B). The result showed similar saccharification efficiency to what was earlier reported when the secretome of *Pf*Mig1^88^ was used for saccharification of acid treated sugarcane bagasse and rice straw (26). The remarkable increase in biomass hydrolyzing capacity of *Pf*Mig1^88^ secretome could majorly be attributed to an increased proportion of CBH, EG and BGL being secreted due to the absence of functional CCR in *Pf*Mig1^88^ as recently reported (26). The 64% hydrolysis achieved with the secretome of *Pf*Mig1^88^ at 20% solid loading was highly significant as it has previously been reported that most saccharification experiments carried out at high substrate loading may not proceed beyond 60% even after physical optimization of the saccharification process (48–50). Also, performing hydrolysis at high substrate loading usually require the addition of high concentration of enzymes which ultimately will not make the process cost-effective. Therefore, efforts have been tailored towards optimizing the cellulase system of fungal strains by fortifying the individual cellulase component in the secretome or supplementing the cellulase with some accessory enzymes such as LPMO, hydrophobins and swollenin (15, 48).

**Fig. 11:**
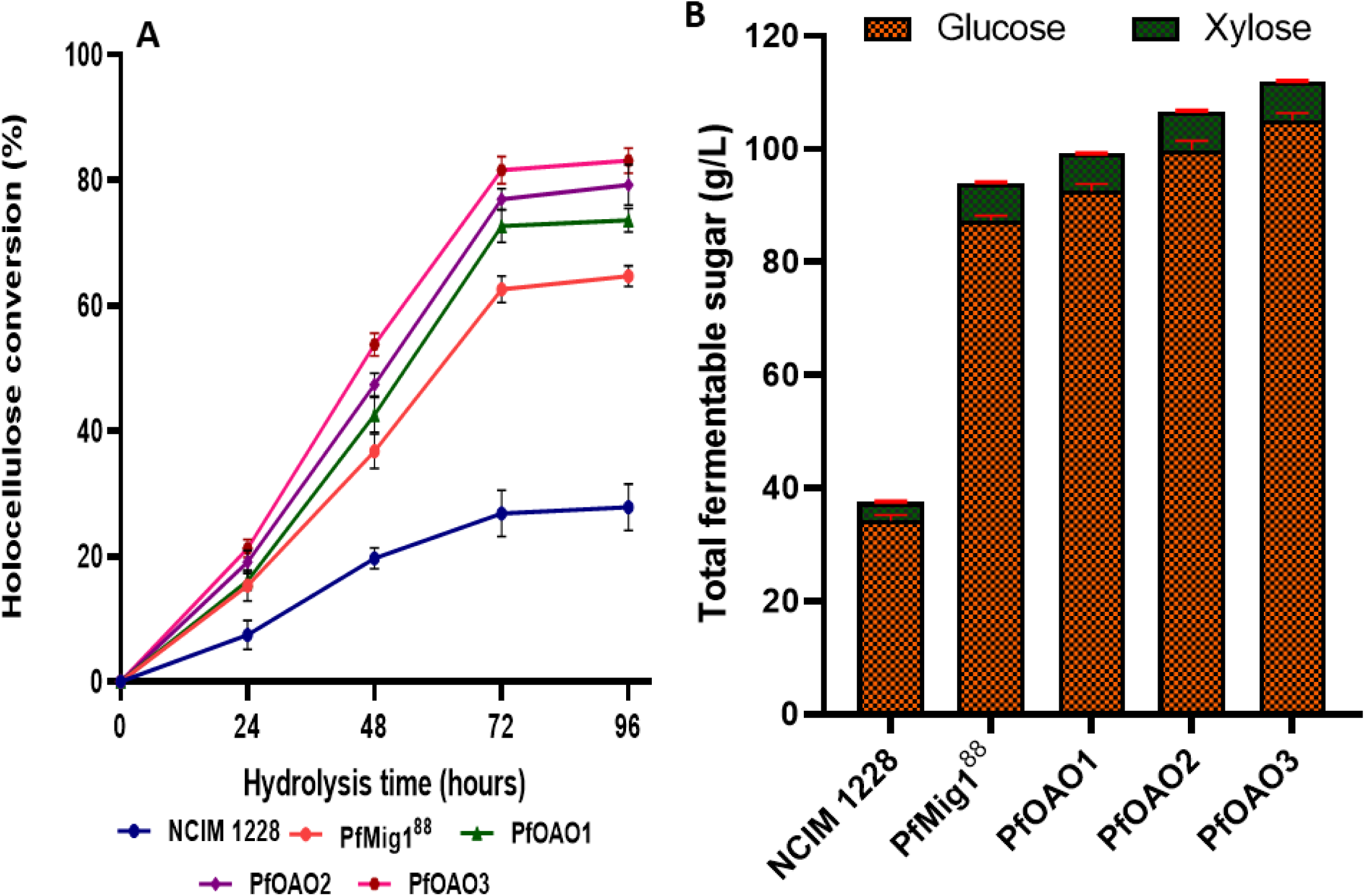
Time course saccharification of nitric acid pretreated wheat straw by the secretome of NCIM1228, *Pf*Mig1^88^ and the all engineered strains at 20% solid loading and protein concentration of 30 mg/g biomass. (A) Percentage sugar release measured at 24 h interval over 96 h saccharification period, (B) total fermentable sugar obtained at 72 h saccharification time point.

Consequently, when the hydrolysis ability of the cellulases produced by the LPMO overexpressing *Pf*OAO1 strain on PWS relative to *Pf*Mig1^88^ was evaluated, a marginal increase in the concentration of monomeric sugars and cellulose conversion was observed (Fig. 11A, 11B). This is because the LPMO alone cannot hydrolyze cellulose on its own and will thus depend on the action of the other hydrolytic enzymes especially CBH1 for optimum saccharification to be achieved as recently reported (48, 51). The total monomeric sugar released by secretome of CBH1 overexpressing *Pf*OAO2 strain was 107 g/L, which corresponded to 77% holocellulose conversion and a 12% increase over that of *Pf*Mig1^88^. The enhanced saccharification obtained with the *Pf*OAO2 strain could not only be due to the increase in cellobiohydrolase level in the system but also linked to the enhancement in the production of β-glucosidase as seen earlier (Fig. 10D). This must have facilitated the enhanced enzymatic degradation of PWS since BGL has the capacity to hydrolyze cellobiose to glucose in the final step and relieve the feedback inhibition of cellobiose on the activities of CBH and EG (52). As expected, there was no significant increase in the hydrolysis of hemicellulose in all the engineered strains when compared with that of *Pf*Mig1^88^ (Table S3).

Interestingly, when both LPMO and CBH1 were co-overexpressed, the total reducing sugar released by the secretome of the strain increased to 112 g/L corresponding to 82% holocellulose conversion (Fig. 11A, 11B). This shows that the simultaneous overexpression of LPMO and CBH1 provided a highly significant increase in saccharification efficiency compared to NCIM1228 (∼199% increase), *Pf*Mig1^88^ (∼20% increase) and the individual ones (∼13% and ∼6% increase for LPMO and CBH1 overexpressing strains, respectively). The results, therefore, suggest that co-overexpression of major cellulase components and its accessory enzyme could facilitate the creation of a more efficient cellulolytic system for optimal hydrolysis of cellulosic biomass at high substrate loading (51, 52).

In conclusion, multiple fungal genetic tools were utilized to construct a more versatile cellulase system for improving the saccharification performance of a catabolite derepressed strain of *P. funiculosum* via overexpression of key oxidative and hydrolytic enzymes in its genome. Combining overexpression of LPMO and CBH1 provided a significantly more efficient cellulase cocktail with an enhanced saccharification performance on PWS at high substrate loading. The LPMO/CBH1 engineered cellulase system exhibited a significantly enhanced cellulose conversion after 72 h enzymatic saccharification of PWS at 20% loading over that of *Pf*Mig1^88^. Results from this study showed that the engineered strains developed could potentially be used as promising bioresource needed for the production of richer and more balanced cellulase cocktail required for low-cost production of lignocellulose-based biofuels.

## Materials and Methods

### Plasmids and Microbial strains

All the strains and plasmids used in this study are listed in Table 5 while the nucleotide sequences for LPMO and CBH1 for the study are stated in the Supplementary file S1. *Escherichia coli* DH5α was used for plasmid propagation throughout the experiments. *Agrobacterium tumefaciens* LBA4404 strain used for fungal transformation was maintained on low sodium LB medium (10 g/L tryptone 5 g/L yeast extract, 5 g/L sodium chloride) containing 100 µg/mL kanamycin and 30 µg/mL rifampicin. pBIF vector (53), which was used as backbone vector for fungal transformation, contains hygromycin and kanamycin resistance genes as selective markers for selection of transformants. The pBluescript (pBSK+) vector containing the ampicillin resistance gene was used as a shuttle vector for simultaneous cloning of genes for LPMO and CBH1. *Penicillium funiculosum* NCIM1228 and its derivative *P. funiculosum* Mig1^88^, the fungal strains used for this study, were routinely cultivated on Petri dishes containing low malt extract-peptone (LMP) agar for about 14 days until full sporulation. For selection and maintenance of fungal transformants, LMP medium supplemented with Hygromycin B at 100 µg/mL was used.

**Table 5:** List of plasmids and strains used in the study.

| Strains | Description | References / sources |
| --- | --- | --- |
| *Escherichia coli* DH5α | F- Φ80lacZΔM15 Δ(lacZYA-argF) U169 recA1 endA1 hsdR17 | Invitrogen |
| *Agrobacterium tumefaciens* LBA4404 | Ach5 with pAch5, Δtra Δocc ΔT<sub>L</sub> ΔT<sub>R</sub> Rif<sup>r</sup> | (25) |
| *Penicillium funiculosum* NCIM1228 | Cellulase producing fungus obtained from NCIM (National Culture Collection Centre, Pune, India) | (19) |
| PfMig1<sup>88</sup> | Catabolite derepressed strain of *Penicillium funiculosum* NCIM1228 | (25) |
| PfOAO1 | LPMO overexpression in PfMig1<sup>88</sup> with the help of pOAO1 vector | This study |
| PfOAO2 | CBH1 overexpression in PfMig1<sup>88</sup> with the help of pOAO2 vector | This study |
| PfOAO3 | LPMO/CBH1 co-overexpression in PfMig1<sup>88</sup> with the help of pOAO5 vector | This study |
| **Plasmids** |  |  |
| pBIF | Kanamycin resistant vector containing EGFP | (53) |
| pBSK+ | Standard cloning vector containing the ampicillin-resistant gene | XB-VEC-1221267 |
| pOAO1 | pBIF vector containing the LPMO gene along with its native promoter and terminator | This study |
| pOAO2 | pBIF vector containing CBH1 gene along with its native promoter and terminator | This study |
| pOAO3 | pBluescriptSK(+) vector containing the LPMO gene along with its native promoter and terminator | This study |
| pOAO4 | pOAO3 vector containing CBH1 gene along with its native promoter and terminator | This study |
| pOAO5 | pBIF vector containing LPMO and CBH1 gene along with with their native promoter and terminator | This study |

### Cellulosic substrate and pretreatment

Nitric acid-treated wheat straw was used as the cellulosic substrate in this study. Wheat straw biomass was comminuted in a cutting mill and sieved through 1.5 mm mesh to get uniform size before chemical pretreatment. Pretreatment was carried out using 0.5% nitric acid in a reactor for 30 mins at 120 °C and 18 bars. The pretreated straw was then repeatedly washed until the pH becomes neutral (54). Compositional analysis conducted according to the NREL/TP510-42618 procedure (55) on the pretreated wheat straw yielded a cellulose content of 61.3%, 6.1% hemicellulose, 15.6% lignin, as well as 6% ashes.

### Computational analysis of LPMO from *P. funiculosum* (PfLPMO9)

The LPMO sequence was retrieved from the draft genome sequence of *P. funiculosum* NCIM1228 available in our laboratory (Supplementary file S1) (19). The amino acid sequences of biochemically characterized LPMOs from other species were retrieved from the NCBI database, and molecular phylogenetic analysis by Maximum Likelihood method and Tamura-Nei model was conducted using MEGA-X software. Multiple sequence alignment of PfLPMO9 with C1/C4 oxidizing LPMOs obtained from the phylogenetic analysis was performed using the Clustal Omega multiple sequence alignment program (56). The alignment was visualized using ESPript 3.0 software (57). To build the structural model of PfLPMO9, the LPMO sequence was first compared with all the available PDB structures using psi blast (56). The top six protein structure from the database namely 6HA5 (43), 2YET (58), 6H1Z (43), 2VTC (59), 5ACF (36), 5NLT (60) were selected as a reference for modelling. The LPMO structure was modelled using Modeller version 9.17. For molecular docking of PfLPMO9 and 2YET protein structures with cellohexaose as the ligand, Autodock vina software version 1.12 was used (61) while UCSF Chimera software version 1.14rc was used for molecular graphics and visualization of the protein structures or complex (62).

### Engineering *Penicillium funiculosum* Mig1^88^ for overexpression of LPMO and CBH1

All vectors for fungal expression described in this study were constructed on the backbone of pBIF vector that was earlier constructed using the binary vector pCAMBIA1300 as a backbone (53). Binary vectors for overexpression of LPMO and CBH1 genes from *P. funiculosum* were constructed as follows: The endogenous gene coding for LPMO was amplified from the genome of *P. funiculosum* NCIM1228 using the primers PfLPMO-F and PfLPMO-R (Table S4). The primers were designed according to LPMO sequence containing both its native promoter and terminator which was obtained from the draft genome sequence of the strain. The PCR product obtained was digested with *Sbf*I and *Spe*I restriction enzymes before ligating unto pBIF vector earlier digested with *Pst*I and *Spe*I enzymes to generate the pBIF/LPMO (pOAO1) vector. For overexpression of CBH1, the CBH1 gene (21) was amplified from the genome of the *P. funiculosum* NCIM1228 using the primers PfCBH1-F1 and PfCBH1-R1 spanning its native promoter and terminator (Supplementary file S1). The PCR product containing *Sac*I and *Bam*HI restriction sites was digested and ligated unto pBIF at the corresponding sites to create the pBIF/CBH1 (pOAO2) vector. The ligated products of pOAO1 and pOAO2 were then transformed in *E. coli* DH5α cells and selected on 50 µg/ml kanamycin. Resulting colonies were screened for positive transformants by colony PCR, followed by restriction digestion of the corresponding plasmids. To create a strain for double overexpression of both LPMO and CBH1 genes, sequential cloning of the two genes was adopted using pBluescript (pBSK+) as a shuttle vector. First, the LPMO gene which had earlier been amplified from the genome was cloned into the pBSK vector between the *Pst*I and *Spe*I site of its multiple cloning sites (MCS) to obtain the plasmid pBSK/LPMO (pOAO3). Secondly, CBH1 gene with its native promoter and terminator was further amplified from the genome of the *P. funiculosum* NCIM1228 using the primers PfCBH1-F2 and PfCBH1-R2 containing *Afl*II and *Spe*I restriction sites, respectively. The PCR product obtained was digested with *Afl*II and *Spe*I restriction enzymes before ligating at the corresponding sites of pOAO3 vector to obtain pOAO4 vector. The resulting *Sbf*I/*Spe*I fragment obtained in the pOAO4 plasmid was excised and cloned between the *Pst*I/*Spe*I sites of pBIF vector, thereby generating the pBIF/LPMO/CBH1 (pOAO5) vector. Positive plasmids of pOAO1, pOAO2 and pOAO5 were transformed to *Pf*Mig1^88^ fungal strain following the Agrobacterium-mediated fungal transformation method (AMTM) as earlier described (63).

### Molecular analysis of transformants

After transformation, the resulting hygromycin resistant transformants were screened following the method described by Fang and Xia (64). The transformants were verified both by PCR and Southern blot hybridization for the integration of LPMO and CBH1 gene expression cassette. For PCR analysis, the primers PgpdA-F and TrpC-R were used to screen for LPMO and CBH1 integration while transformants of LPMO/CBH1 were screened using PgpdA IR-F and PfCBH1-IR R primer set (Table S4). Southern hybridization was performed using standard procedures described by Southern, 2006 (65). Genomic DNAs (8 µg) of *Pf*Mig1^88^ and transformants of LPMO, CBH1 and LPMO/CBH1 were digested with *Xho*I, *Hind*III and *Apa*I restriction enzymes. The digested gDNAs were then size-fractionated by electrophoresis on 0.8% agarose gel in 1× TAE (Tris-acetate) buffer. After depurination using 250 mM HCl, denaturation (1.5 M NaCl and 0.5 M NaOH) and neutralization (1.5 M NaCl and 1.0 M Tris–HCl pH 8.0) steps, the gel was capillary-blot onto a positively charged Hybond™-N^+^ membranes (Amersham Biosciences, USA). The 621-bp fragment of LPMO gene was PCR-amplified with the primers LPMO-Probe F and LPMO-Probe R and labelled as a probe to detect the LPMO integration. Similarly, a fragment of CBH1 gene (605-bp long) was amplified using the primers CBH1-Probe F and CBH1-Probe R and used to confirm the insertion of the CBH1 expression cassette in both CBH1 and LPMO/CBH1 strains (Table S4). The LPMO and CBH1 amplicons for detection were radio-labelled with [α-32P] dCTP using the NEBlot Kit (NEB, USA) according to the manufacturer’s instructions and used as a probe to determine the copy number of T-DNA integrations in all the transformants.

### Expression analysis of cellulolytic enzymes via real-time PCR

For real-time PCR experiments, cultures of all strains were grown in Minimal Mandel’s media containing 4% Avicel for 48 h (66). Mycelia were harvested by filtration and frozen in liquid nitrogen. RNA was extracted using RNeasy kit (Qiagen) according to the manufacturer’s instructions. RNA was DNase treated (Invitrogen) before cDNA synthesis. One microgram of RNA was used as template in each quantitative real-time PCR (qRT-PCR) reaction. cDNA synthesis control was performed to ensure the absence of DNA contamination. qRT-PCR was carried out using iTaq™ Universal SYBR® Green Supermix (Bio-Rad) and Bio-Rad CFX96 qPCR detection system. Primers for transcripts to be tested were designed using boundary sequence of two exons to avoid any amplification from genomic DNA contamination. qRT-PCR was done in biological triplicates with tubulin as the endogenous control. Relative expression levels were normalized to tubulin, and fold changes in RNA level were the ratios of the relative expression level of *Pf*Mig1^88^ and the corresponding transformants of LPMO and CBH1 to NCIM1228 under cellulase inducing conditions (67).

### Cellulolytic secretome preparation

*Penicillium funiculosum* NCIM1228, *P. funiculosum* Mig1^88^ (*Pf*Mig1^88^) and the resulting transformants of LPMO, CBH1 and LPMO/CBH1 were cultivated on Petri dishes containing low malt extract agar until there was full sporulation. After 14 days of incubation, spores were recovered with sterile water, filtered through sterile Mira cloth and quantified using a hemocytometer. The primary culture of each strain was prepared by culturing 10^7^ conidiophores in potato dextrose broth (PDB) for 36 h. Primary cultures of the strains were added to the cellulase inducing media in Erlenmeyer flasks at a final concentration of 10%. Cellulase inducing media contains soya peptone (24 g/l), wheat bran (21.4 g/l), microcrystalline cellulose (MCC) (24 g/l), KH_2_PO_4_ (12.4 g/L), K_2_HPO_4_ (2.68 g/l) (NH4)2SO_4_ (0.28 g/l), CaCO_3_ (2.5 g/l), corn steep liquor (1%), urea (0.52 g/l), and yeast extract (0.05 g/l) with the final pH adjusted to 5.0. The flasks were kept at 28 °C for five days with orbital shaking at 150 rpm (Innova 44, Eppendorf AG, Germany). Induced cultures were centrifuged at 9000 rpm for 10 min at 4 °C, and the cellulolytic supernatants were collected and stored at 4 °C until use.

### Determination of enzyme activities of the engineered secretome

All enzymatic activities performed in this study were routinely determined following standard assay procedures. Cellobiohydrolase activity was determined by incubating appropriate dilution of the enzyme with 1% Avicel^®^ PH-101 (Sigma) for 120 mins. Endoglucanase, xylanase and β-glucosidase activities were determined by incubating appropriate dilution of the enzyme with 2% CMC (Sigma), 2% beechwood xylan (HiMedia) and p-nitrophenyl-β-D-glucopyranoside (Sigma), respectively, for 30 mins, after which the amount of reducing sugars released was measured as previously reported (25). One unit of CMCase, Avicelase and xylanase activity is defined as the amount of enzyme releasing 1 μmol of reducing sugar per min while one unit of β-glucosidase activity was defined as the amount of protein that released 1 μmol of p-nitrophenol per min. LPMO activity was assayed using the Amplex red assay, as described earlier (16). The reaction mixture was composed of 20 μL of LPMO source (enzyme) and 180 μL assay solution, which comprised 18 μL of 300 μM ascorbate, 18 μL of 500 μM Amplex Red, 18 μL of 71.4 units/mL Horse Radish Peroxidase (HRP), 18 μL of 1 M sodium phosphate buffer pH 6.0, and 108 μL of HPLC-grade water. Resorufin fluorescence was taken at an excitation wavelength of 530 nm and emission wavelength 580 nm after 10 min of incubation at 22 °C using a multimode plate reader (Spectra Max M3, USA). In reference experiments without LPMO, the background signal was measured and subtracted from the assays. A standard curve obtained with various dilutions of H_2_O_2_ was used for the calculation of an enzyme factor to convert the fluorimeter readout (counts min^−1^) into enzyme activity. LPMO activity is defined as 1 μmol of H_2_O_2_ generated per minute under the defined assay conditions. Total cellulase activity in the secretome was measured in terms of filter paper units (FPU) per millilitre of original (undiluted) enzyme solution. The assay requires a fixed degree of conversion of substrate, from 50 mg of filter paper within 60 min at 50 °C. The FPU is defined as the amount of enzyme required to produce 2 mg of glucose from 50 mg of filter paper within 60 min of incubation. Total protein of each secretome was estimated by the Bicinchoninic acid (BCA) using bovine serum albumin (BSA) as standard.

### SDS-PAGE and Zymogram Analysis

Sodium dodecyl sulfate (SDS)-polyacrylamide gels (12%) were prepared, and proteins obtained following the culture supernatant preparation were separated via SDS-polyacrylamide gel electrophoresis (PAGE) according to method earlier described (68). Mini-PROTEAN Tetra Cell (Bio-Rad) with gel size of 8.6 × 6.7 cm^2^ was used for protein separation. After protein separation, the gels were first washed with 1x PBS buffer to remove the SDS. The Zymogram analysis was carried out using 5 mM 4-methyl umbelliferyl β-D-glucopyranoside (MUG) as substrate following standard procedures in 50 mM citrate-phosphate buffer pH 4.0 for 10 mins before visualization under UV light. Proteins of the gel were then stained with Coomassie blue R-250 (Sigma-Aldrich, USA) and the molecular mass of the proteins was determined with reference to standard proteins (Thermo Scientific, USA).

### Biomass saccharification and quantification of fermentable sugars

The saccharification efficiency of the secretome of all the strains used in the study towards pretreated wheat straw was carried out according to the method described by Ogunmolu et al. (19) with some modifications. Performance of the secretomes towards nitric acid-treated wheat straw was evaluated at 20% dry weight of biomass using an enzyme concentration of 30 mg/g biomass. Saccharification was performed in 50 ml screw-capped Falcon tubes in an incubator shaker at 50 °C for 96 h. The reaction mixture included the nitric acid-treated wheat straw at 20% dry weight loading in a 5 ml final reaction volume. Total protein content in the secretome of all the fungal strains tested was measured, and the appropriate volume of desired protein concentration (30 mg/g_DBW_) was added to the reaction mixture. The reactions were set up in 100 mM citrate-phosphate buffer (pH 4.0) and incubated at 50 °C with constant shaking at 300 rpm for 96 h. Samples were collected every 24 h and analyzed for production of fermentable sugars. Control experiments were carried out under the same conditions using substrates without enzymes (enzyme blank) and enzymes without substrates (substrate blank); a substrate-free negative control was set up by filling the Falcon tubes with 100 mM citrate-phosphate buffer, pH 4.0, and the background of soluble sugars present in the respective biomass was determined by incubating each biomass in the absence of enzyme. Following the completion of hydrolysis at each time point, the Falcon tubes were centrifuged at 3500 rpm for 10 min in a swinging bucket centrifuge (Eppendorf, Germany) to separate the solid residue from the digested biomass. Supernatants which were recovered after enzymatic hydrolysis of the pretreated wheat straw were analyzed by high-performance liquid chromatography equipped with Aminex HPX-87H anion exchange column (Bio-Rad, USA) and a refractive index detector to analyze released monosaccharides (glucose and xylose) by anion exchange chromatography. The filtered mobile phase (4 mM H_2_SO_4_) was used at a constant rate of 0.3 ml/min with the column, and RI detector temperatures maintained at 35 °C. The concentration of each monosaccharide was calculated from calibration curves of external standards (xylose and glucose) purchased from Absolute Standards Inc. The following equations provided in NREL’s LAP TP-510-43630 was then used to determine the theoretical conversions of cellulose and hemicellulose (in percentage) into monomeric sugars;

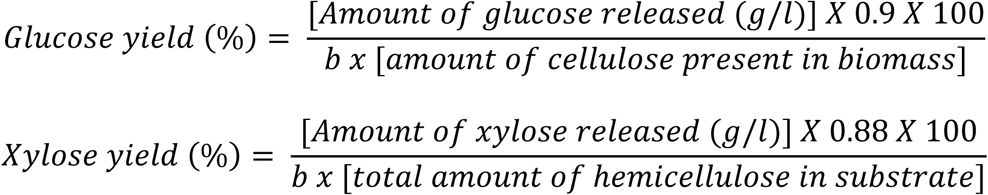

Where ***b*** is the cellulose or hemicellulose fraction in the dry pretreated biomass (g/g) (69, 70).

### Data and statistical analysis

All experiments were performed in triplicate, and results are presented as mean and standard deviation. The data was collected in a Microsoft Excel spreadsheet where the average and standard error of the mean were determined. All graphs were created using the GraphPad Prism 8.0 software. Data were further evaluated by one-way analysis of variance (ANOVA) and multiple t-tests using GraphPad Prism 8.0 software where appropriate.

## Supplementary information

E-supplementary data of this work can be found in the online version of the paper

## Acknowledgements

This study was funded by Department of Biotechnology, Government of India via Bioenergy Centre grant no. BT/PR/Centre/03/2011. The ICGEB Fellowship Unit is highly appreciated for the Arturo Falaschi Pre-Doctoral Fellowship awarded to OAO. Authors would like to thank Prof. Arvind M. Lali for providing pretreated biomass used for this study.

## Supplementary information details -

**Fig. S1: Structural model of PfLPMO9.** Panel A represents the best model of the PfLPMO9 catalytic modules with the amino acids interacting with copper ions. The loop regions that potentially contribute to functional variation among LPMOs, named L2, L3, LS and LC are marked with dark blue, light blue, green and purple, respectively. Panel B is the Ramachandran plot validation of the modelled structure evaluated by PROCHECK.

**Fig. S2: 3-D structures of the top three models of PfLPMO9 and TaLPMO9A (2YET) bounded to celllohexaose** with their corresponding binding energies

**Fig. S3: Molecular confirmation of recombinant vectors and engineered strains** (a) Restriction digestion analysis of pBIF (1) and pOAO1 (2) by *Bam*HI. (b) Determination of PfLPMO9 integration in the genome of *Pf*Mig1^88^ transformants by PCR using the primers PgpdA-F and TrpC-R. The expected size is 4.8 kbp, corresponding to a fragment spanning the region of gpdA promoter and TrpC on the expression cassette. Lane M is DNA molecular mass marker; lane (+) is the pOAO1 plasmid which served as a positive control, lanes 1–10 are the transformants of pOAO1, lane (−) is the gDNA of *Pf*Mig1^88^ which was the negative control. (c) Restriction digestion analysis of pBIF (1) and pOAO5 (2) by *Afl*II and *Spe*I. (d) Determination of LPMO/CBH1 cassette integration in the genome of *Pf*Mig1^88^ transformants by PCR using the primers PgpdA IR-F and PfCBH1 IR-R. The expected size is 5.0 kbp, corresponding to the T-DNA region, which contains the fused fragment of LPMO and CBH1 on the expression cassette. Lane M is DNA molecular mass marker; lane (+) is the pOAO5 plasmid which served as a positive control, lanes 1–6 are the transformants of pOAO3, lane (−) is the gDNA of *Pf*Mig1^88^ which was the negative control.

**Fig. S4: Construction of CBH1 expression cassette and its overexpression in *Pf*Mig1**^**88**^.

(a) Schematic diagram showing the assembly of CBH1 cassette from *P. funiculosum* NCIM1228 genome. (b) Restriction digestion analysis of pBIF (1) and pOAO2 (2) by *Sac*I and *Bam*HI. (c) Transformants of pOAO2 after AMTM transformation in *Pf*Mig1^88^. Transformants were selected on 100 µg/ml hygromycin. (d) Determination of PfCBH1 integration in the genome of *Pf*Mig1^88^ transformants by PCR using the primers PgpdA-F and TrpC-R. The expected size is 5.2 kbp, corresponding to a fragment spanning the region of gpdA promoter and TrpC on the expression cassette. Lane M is DNA molecular mass marker; lane (+) is the pOAOA2 plasmid which served as a positive control, lane 1–8 are the transformants of pOAOA2, lane (−) is the gDNA of *Pf*Mig1^88^ which was the negative control. (e) Southern blot of transformants confirmed by PCR. M is *Hind*III lambda DNA size marker used; Lane C indicates genomic DNA of non-transformed parental strain; Lanes 1-6 represent genomic DNA from transformants with CBH1 cassette integration; (f) The cellobiohydrolase and FPase activities in the fermentation broth of *Pf*Mig1^88^ and six transformants.

**Fig. S5**: **Protein profile and zymogram assay for identification of β-glucosidase in the secretome of all strains.** Culture supernatant from all strains were separated in 10% SDS-PAGE. β-glucosidase activity was detected by MUG-zymogram assay. Lane M, molecular weight marker. Lanes 1, 2, 3, 4, 5 were secretome of NCIM 1228, *Pf*Mig1^88^, *Pf*OAO1, *Pf*OAO2 and *Pf*OAO3, respectively.

**Table S1:** Comparative evaluation of the molecular interactions of PfLPMO9, TaLPMO9A and LsLPMO9A bounded with cellohexaose

**Table S2:** Specific Activities (U/mg protein) of selected biomass hydrolyzing enzymes of *P. funiculosum* NCIM1228, *Pf*Mig1^88^ and its resulting transformants

**Table S3:** yield of fermentable sugars obtained at 72h saccharification of nitric acid treated wheat straw by the secretome of NCIM1228, *Pf*Mig1^88^ and all engineered strains

**Table S4:** List of primers used in this study

**Supplementary File 1:** Construction of CBH1 strain, and nucleotide sequence of PfLPMO and PfCBH1.

